# The Metabolome of *Drosophila* Reveals Exercise-Induced Stress and Anti-Inflammatory Response

**DOI:** 10.64898/2026.06.26.734830

**Authors:** Tolulope R. Mabayoje, Anne E. Backlund, Jordan E. Albrecht, Mckenzie S. Chamberlain, Sean S. Tremblay, Abigail Myers, Alyssa Koehler, Miled A. Maisonet-Nieves, Cross Hogland, Gabriella Bicanovsky, Jarmacus Monroe, Laura K. Reed

## Abstract

Exercise is a known remedy for metabolic disorders, including obesity and type II diabetes. In humans and some model organisms, exercise is classified into different types and intensities based on the mode of exercise and the devices used. Since *Drosophila* has emerged as a model for exercise research, several devices have been developed for exercise training. These devices vary in mechanisms of operation; for example, the Power Tower (PT) uses a repeated drop-and-hit mechanism, and the TreadWheel (TW) uses end-over-end rotation. This variation in mechanisms that trigger the negative geotaxis of flies could impact exercise outcomes, and it remains unclear how this variation drives the response of flies to exercise training. Stress is a known response to exercise in humans and mice. Stress in Drosophila can be induced by various conditions, including heat, cold, chemical exposure, and other environmental stressors, but there is little information on exercise-induced stress. It is also unclear whether different devices elicit distinct stress responses or whether the exercise-induced stress response is similar to other stress responses (e.g., thermal, chemical). To explore the effects of device, sex, and genotype on multiple metabolic pathways, we used an untargeted metabolomics approach to further elucidate the molecular mechanisms of exercise. As no studies have directly examined the exercise metabolome of *Drosophila*, this study is the first to explore the metabolome of flies post-exercise. Our results revealed that exercising on different devices elicits varying responses and reshapes the metabolome of flies. We also identified specific small compounds that may be linked to stress and inflammatory response more in the PT-exercised flies than in the TW-exercised flies. Overall, the exercise device effect interacted with sex effect and genotype, such that male and female flies have different metabolite compositions post-exercise on TW and PT.

## Introduction

Exercise is a lifestyle recommendation and non-invasive treatment option that is known to mitigate metabolic syndrome, which includes obesity, diabetes, cardiovascular disease, and other metabolic diseases (Higuera-Hernández et al., 2018; Ruegsegger & Booth, 2018; Ozemek et al., 2020). Exercise also has the potential to prevent neurodegenerative diseases such as Alzheimer’s and Parkinson’s (Liu et al., 2019; Sujkowski et al., 2022). Model organisms (e.g., mice, *Drosophila*, and *C. elegans*) are rich resources that have provided helpful insight into the potential benefits of exercise (Xu et al., 2017; Sujkowski & Wessells, 2018; Pham et al., 2021; Furrer et al., 2023). For example, endurance exercise in mice and swimming exercise in *C. elegans* delayed the onset of neurodegenerative diseases (Xu et al., 2017; Bhadra et al., 2023).

The beneficial effects of exercise in *Drosophila* were first established in 2009 with the Power Tower (PT) exercise device (Piazza et al., 2009). Since the PT emerged as an exercise training device, other researchers have developed various exercise devices, including the TreadWheel (TW) (Mendez et al., 2016), the Rotating Exercise Quantification System (Watanabe & Riddle, 2017), and the horizontal steel flipping tubes (Zheng et al., 2015). All these devices rely on negative geotaxis to induce exercise; however, the mechanisms that induce flies’ negative geotaxis vary across devices (e.g., PT uses a drop-and-hit mechanism, while the TW uses an end-over-end rotation). This variation in mechanism could affect flies’ response to exercise training. Therefore, to characterize this device effect first, we compared flies trained on the PT and TW across multiple genotypes.

Among factors that contribute to the effectiveness of exercise, sex and genetic variation play crucial roles. Sex-based differences in fuel utilization and hormonal environment are well-documented modifiers of exercise response in vertebrates and flies (Friedlander et al., 1998; Hunter, 2014; Abo et al., 2024). In the context of endurance exercise in humans and flies, females have shown a greater metabolic advantage when trained at the same submaximal intensity (Hunter, 2014). There is also a sex difference in the relative utilization of carbohydrates and lipids as fuel sources in humans, both at rest and during exercise (Abo et al., 2024). These studies have revealed that differences in respiratory rate between men and women disappear during high-intensity exercise and when individuals are well adapted to exercise (Friedlander et al., 1998; Abo et al., 2024). Also, in *Drosophila*, many studies have reported genotype and sex-specific responses to exercise (Backlund et al., 2025; Johnson & Riddle, 2024; Watanabe et al., 2020; Mendez et al., 2016). Furthermore, a Genome-Wide Association Study (GWAS) classified flies into marathon and sprint runners based on underlying genetic differences (Watanabe et al., 2020). However, the interaction between genetic variation and exercise device on exercise adaptations has not been studied.

In this study, we compared exercise responses across multiple genotypes to characterize how genetic variation shapes exercise outcomes across devices. We assessed variation in fitness by climbing performance (see Methods) in males and females across six genetic lines obtained from the *Drosophila* Genetic Reference Panel (DGRP) (Mackay et al., 2012).

One of the benefits of exercise is increased mitochondrial function and biogenesis in humans and mice (Garanata et al., 2018; Memme et al., 2021). This increase in mitochondrial activity produces free radicals and reactive oxygen species (ROS), which increase stress (Chen et al., 2024), yet exercise is also linked to antioxidant and anti-inflammatory effects (El Assar et al., 2022; Kawanishi et al., 2013). Since exercise can both increase mitochondrial ROS production and mitigate the detrimental effects of free radical production caused by oxidative stress, it is unclear how exercise shifts from an antioxidant effect to promoting ROS production and how different exercise regimens contribute to this shift. Additionally, ROS accumulation causes inflammation in humans (Steensberg et al., 2000), and exercise has an anti-inflammatory effect in mice and humans (Petersen & Pedersen, 2005; Scheffer & Latini, 2020). It remains unclear which stress responses exercise induces in *Drosophila,* and whether exercise could combat inflammation while simultaneously enhancing stress responses. Although a level of exercise training is necessary to improve muscle strength and induce exercise adaptation, no thresholds have been established for the exercise intensities that cause stress in flies (Sharma & Mehdi, 2023).

In *Drosophila*, stress can be induced through chemical exposure, thermal challenge, or other environmental stressors. Chemicals commonly used to model oxidative stress include paraquat, bisphenol, and cadmium, while thermal stress is induced through heat or cold exposure (Hosamani & Muralidhara, 2013; Colinet et al., 2010; MacMillan et al., 2016; Chen et al., 2022; Yang et al., 2022; Ramalho et al., 2024). These conditions elevate established biomarkers of oxidative damage, including malondialdehyde, hydroperoxide, and superoxide dismutase activity. However, exercise-induced stress patterns have not been thoroughly studied in *Drosophila.* Since exercise-induced stress in *Drosophila* is poorly characterized, we exercised flies on the PT and TW to identify device-specific stress signatures. We hypothesized that the sudden deceleration of flies during exercise on the PT device would induce greater physiological stress than the TW’s gentler rotational mechanism.

Our study compared the expression patterns of *heat shock protein* 22 (*Hsp22*), which is upregulated by heat, oxidative, and cold stress in *Drosophila* (Colinet et al., 2010; Morrow et al., 2016), across exercise treatments. Other genes we studied include *gag-related protein* (*Gagr*) and *unpaired 3* (*Upd3*), which can be upregulated when the JAK/STAT pathway is activated (Gigin & Nefedova, 2023). The JAK/STAT pathway is known to regulate stress response and innate immunity in *Drosophila* (Nefedova et al., 2022; Gigin et al., 2023). We also studied the expression of genes associated with mitochondrial biogenesis: *spargel (srl*) and *dynamin-related protein-1* (*drp-1*) (Chang & Blackstone, 2010; Tiefenbock et al., 2010). These genes regulate mitochondrial biogenesis and mitochondrial fission, respectively (Chang & Blackstone, 2010; Tiefenbock et al., 2010). We measured the expression of these genes across three DGRP lines on both devices to determine whether device type and genotype interact to shape the stress-related transcriptional response to exercise.

Lastly, exercise research in *Drosophila* has aided the identification of genes and pathway changes that contribute to exercise outcomes. Examples include the upregulation of glutathione turnover, downregulation of Akt/p38 MAPK 22, and the upregulation of Nrf2 pathway post-exercise (Dahleh et al., 2023). However, there is little information on the concurrent impact of exercise across multiple pathways. Metabolomic studies can provide insight into the contributions of hundreds to thousands of metabolites. Profiling these compounds simultaneously can resolve how exercise reshapes multiple metabolic pathways and how external factors (device type, sex, genotype) modulate that response. In mice, for example, targeted metabolomics identified exercise-inducible metabolites that suppress obesity (Li et al., 2022).

The *Drosophila* metabolome has been explored to better understand the aging process, the interaction of genotype and environmental factors, and the effects of heat stress (e.g., Malmendal et al., 2006; Reed et al., 2014; Zhao et al., 2022). In the *Drosophila* metabolome, coenzyme A was identified as protective against toxicity induced by a high-sugar diet (Palanker et al., 2016). Dietary restrictions also shift the metabolome, reducing activity in the pentose phosphate pathway, gluconate pathway, and trehalose degradation across abdominal tissues in a lifespan-extending manner (Laye et al., 2015). However, no studies have specifically examined the *Drosophila* metabolome after exercising to uncover the mechanism by which exercise impacts metabolic pathways.

Using an untargeted approach, we explored the *Drosophila* metabolome to identify how different exercise devices, sex, and genotype affect the products of multiple pathways. Characterizing the *Drosophila* exercise metabolome may reveal conserved molecular responses to exercise and establish this system as a tractable model for translational exercise research.

## Materials and methods

This study is comprised of three experiments. Experiment 1 assessed climbing performance across six DGRP genotypes (907, 900, 805, 440, 802, 748, and 440) on the PT and TW following five days of exercise. Experiment 2 measured the gene expression in three genotypes (DGRP 440, 802, and 805) after a single 30-minute exercise session, followed by expression profiling in DGRP 440 after five days of continuous exercise. Experiment 3 used untargeted GC-MS metabolomics to characterize the post-exercise metabolome of two genotypes (DGRP 440 and 900) selected based on the outcomes of Experiments 1 and 2. The exercise protocol for Experiments 1 and 3 is described under Exercise Training below.

### Fly Stocks

We used six *Drosophila* Genetic Reference Panel (DGRP) lines to assess climbing performance: DGRP 907, 900, 805, 802, 748, and 440 (Bloomington stock IDs 28262, 28261, 28237, 28235, 28224, and 28197, respectively). These genetic lines were randomly selected with no prior knowledge of their response to exercise training. All replicates for a given strain were completed before beginning the next strain. To control for population density, we placed the parent stocks in an egg-laying chamber, and 50 first-instar larvae were collected into vials containing a standard cornmeal molasses food (by weight 5.28% cornmeal, 1.05% yeast, 0.56% agar, 87.03% water, 4.37% molasses, 1.15% Tegosept, 0.55% propionic acid). These vials were maintained in the incubator at 25 °C for 10-12 days until the adults emerged. Young flies were collected for four days post-eclosion, sexed, and randomly assigned to treatments (Treadwheel (TW), Power Tower (PT), or no exercise (NE)). We placed ten male or female flies in each vial, and each treatment had three replicates per sex. The three treatment groups for each device were exercise (E; with 6 cm of exercise space), control (C; with only 1 cm of space), and no exercise (NE; which was not placed on the device). The NE group was kept adjacent to the exercise devices so that they experienced the same temperature, humidity, and sound as those on the devices.

The abbreviations for each treatment are represented in Table 1.

**Table 1:** Treatment codes and their respective meanings.

| Treatment Code | Meaning |
| --- | --- |
| PTE (M/F) | Power Tower Exercise (Male/Female) |
| TWE (M/F) | TreadWheel Exercise (Male/Female) |
| PTC (M/F) | Power Tower Control (Male/Female) |
| TWC (M/F) | TreadWheel Control (Male/Female) |
| NE (M/F) | No Exercise (Male/Female) - untrained |

### Exercise Training

For experiments one and three, flies started exercise training on the fifth day post-eclosion, and exercised at the same time each day (11 am – 1 pm) for five consecutive days. The two-hour session consisted of six cycles of 15 minutes of exercise at four rounds per minute (rpm) followed by five minutes of rest. After the fifth day, the flies were placed in the incubator to recover one day before analyzing their climbing performance. In experiment two, flies from three genetic lines (DGRP 440, 802, and 805) were given a single 30-minute exercise session at five days post-eclosion to represent an acute exercise condition. These genotypes were selected because of their distinct performance on the PT and TW (DGRP 440’s climbing performance declined on the TW but improved on the PT, while DGRP 802 and 805 improved on the TW and declined on the PT). Gene expression was then measured in all three genotypes after the 30-minute session and in DGRP 440 following five days of exercise.

### Climbing Performance Assay

The climbing performance assay was conducted one day after the cessation of exercise. We transferred flies from each experimental replicate into a clean, empty vial and placed a piece of clear parafilm over the vial to seal it. We left these flies in the new vial for at least five minutes to acclimate. We then placed one vial at a time in front of a board with a grid scaled in 1 cm squares. We tapped the flies to the bottom of the vial three times, then took a picture of the flies three seconds later with a digital camera placed at a fixed distance. We repeated this process three times for each vial.

We used ImageJ software (Rueden et al., 2017) to convert the individual climbing heights of the flies to a calculated distance in centimeters. This was determined by equating one grid square to 1 cm. This gave us values for each fly within a vial. Images for each vial were captured three separate times per treatment replicate. For analysis, individual fly measurements were nested within vial across the three image captures, with each vial treated as a replicate. We had three vials per treatment and measured climbing performance across six genotypes. We transferred these data point to JMP, confirmed that they were normally distributed, and analyzed the climbing performance of flies with the general linear model:

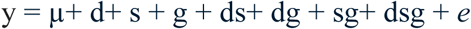

The tested effects were exercise device (d), sex (s), genotypes (g), and their interactions, where ‘e’ and ‘µ’ represent error and population mean, respectively. We also conducted a student’s t-test to compare the climbing performance of unexercised males and females.

### Sample Processing for Gene Expression

For the 30-minute exercise treatment group, ten flies per replicate were frozen in liquid nitrogen immediately after exercise and stored at-80 °C. Three replicates per treatment were collected across all three genotypes (DGRP 440, 802, and 805). DGRP 440 flies from the five-day exercise group were collected by the same method, frozen in liquid nitrogen immediately after the final session, and stored at-80 °C. We extracted RNA using the E.Z.N.A. Total RNA Kit 1 (product no: R6834-02) from Omega BioTek. We used the iScript ^(TM)^ cDNA synthesis kit (product no:1708890) from Bio-Rad Laboratories to convert the RNA to cDNA following the manufacturer’s protocol.

### Gene Expression

To determine the effect of acute exercise training (30 minutes) on gene expression of males and females in three genetic lines (DGRP 805, 802, and 440), we analyzed expression patterns of *spargel* (*srl*), *Gag-Related Protein* (*Gagr*), *Translocator Protein O* (*TSPO*), *heat shock protein 22* (*Hsp22*), *Unpaired 3 protein* (*Upd3*), and *Stress Induced Nuclease* (*sid*). We then compared gene expression patterns after 1 session of 30 minutes to those after five days of exercise training (2 hours per day) in DGRP 440. For this, we considered additional genes: *Dynamin-Related Protein* (*Drp1*) and *Vrille* (*vri*). We used *Ribosomal protein 49* (*Rp49)* as the reference gene for *srl* following (Tinkerhess et al., 2012), and *Ribosomal Protein L 40* (*Rpl40)* as the reference gene for the stress-associated genes (*Gagr*, *Upd3*, *Vrille*, *sid*, *Hsp22),* as it has been validated as a stable housekeeping gene in *Drosophila* (Polymenis, 2020). We obtained the primers from the Fly Primer Bank (Hu et al., 2013; Table 1. S1).

We conducted RT-qPCR using the SsoAdvanced Universal SYBR Green Supermix (catalog no: 1725270) to determine the levels of expression of these genes relative to a pooled sample across our exercise (E) and the no exercise (NE) groups. Following the manufacturer’s procedure, the RT-qPCR master mix consisted of 10 µL of SYBR Green, 1 µL of a 1:20 dilution of 100 mM forward and reverse primers, 3 µL of water per sample, totaling 15 µL. To the 15 µL master mix, we added 5 µL (10 ng/µL) cDNA for each sample in their designated wells. For the pooled sample, we pooled 2 µL of each sample’s final cDNA concentration per genotype into a separate Eppendorf tube. We had three technical replicates of the pooled samples for each RT-qPCR run, and their CT values within the run were averaged for further analysis. To calculate the ΔCT values, we subtracted the *Rpl40 or Rp49* CT values for each run from the CT values of each target gene in each run. We then calculated the ΔCT of our pooled samples by subtracting the average CT values of *Rpl40* or *Rp49* from the average CT value of the pooled samples. The ΔΔCT was calculated by subtracting the average ΔCT value of the pooled samples from the ΔCT values for each replicate.

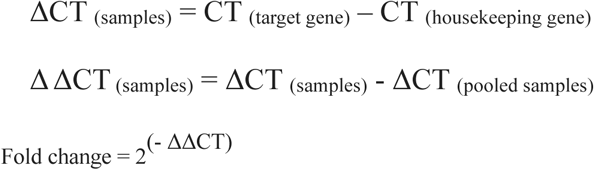

The ΔΔCT and fold change values for each sample were prepared in Excel and exported to JMP. We assessed the normality of our data by visual inspection of histograms and Quantile-quantile (Q-Q) plots prior to analysis. To determine the effects of exercise device, sex, and genotype, we first analyzed gene expression data sets from the 30-minute exercise treatment across three genotypes and followed up with analyses of gene expression data collected from DGRP 440 after a 30-minute and 5-day exercise treatment. Using a generalized linear model, we determined the effects of exercise device, sex, and genotype for individual genes reported in Table 3. To highlight groups with significantly different treatment means, we conducted pairwise comparisons using an unpaired Student’s t-test and represented the data in bar graphs with error bars representing ± SEM (standard error of the mean). The graphs were plotted in PRISM 10 (GraphPad Software, www.graphpad.com).

### Metabolomics

We conducted untargeted gas chromatography mass spectrometry to identify small molecules whose abundance differed between exercised and unexercised flies on the PT and TW. We analyzed the effect of genotype and sex on the metabolome of these flies from two genotypes, DGRP 440 and 900, selected from the six genetic lines previously studied in experiment one (Fig 1). DGRP 440 was selected for its elevated *Hsp22* expression following continuous exercise on both devices (see Results), while DGRP 900 was selected for its improved climbing performance on both devices. We exercised flies using the five-day exercise regime described in experiment 1 (Figure 1.1). At the end of the five days, the flies were frozen in liquid nitrogen immediately after exercising and sent to the UC-Davis Metabolomics facility for untargeted GC-MS analysis. For each replicate, we included 10 flies with three replicates per treatment.

**Figure 1.**
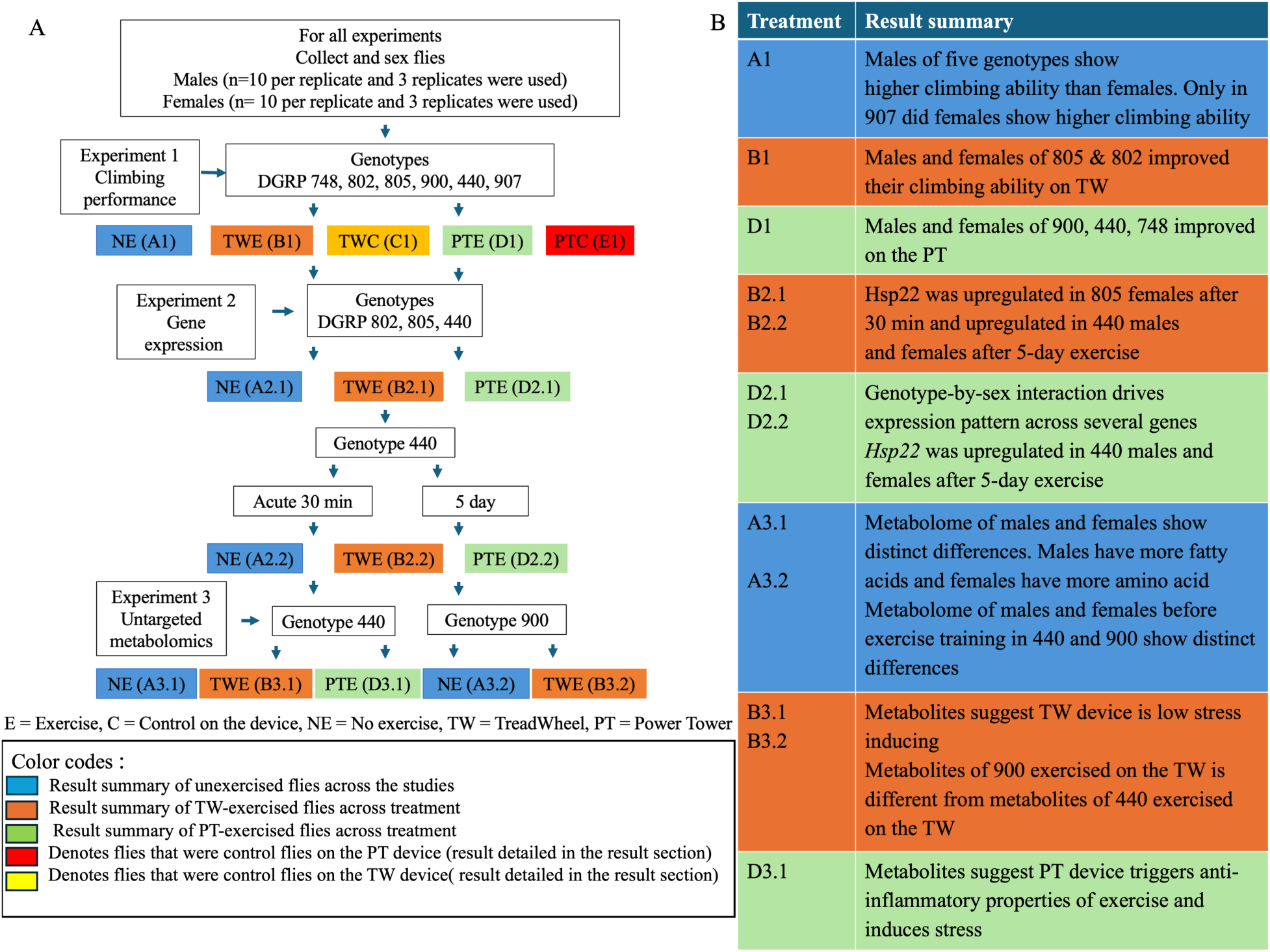
Schematic illustration of our study progression from phenotypic analyses to gene expression and then metabolomics of exercised flies. **A)** The chart shows the progression of experiments one, two, and three. **B)** Represents a summary of the experiment’s results.

### Metabolomics Data Acquisition

The samples were prepared and extracted according to the protocols described in Fiehn et al. (2008). Briefly, the column used was the Restek corporation Rtx-5Sil MS (30 m length x 0.25 mm internal diameter with 0.25 µm film made of 95% dimethyl/ 5% diphenylplysiloxane). Helium was used for the mobile phase, and the column temperature was kept within the range of 50-330 °C. The flow rate was maintained at 1 mL/min, with an injection volume of 0.5 µL. Samples were injected in splitless mode (25 s splitless time). The injection temperature was 50 °C and then increased to 250 °C by 12 °C increment per second. The oven temperature program was set at 50 °C for one min, then using 20 °C increments per minute the temperature was increased to 330 °C, then held constant for 5 minutes. The analytical GC column chromatography method yields retention and separation of primary metabolite classes (amino acids, hydroxyl acids, carbohydrates, sugar acids, sterols, aromatics, nucleosides, amines, and miscellaneous compounds). Metabolites must have narrow peak widths of 2–3 s and very good within-series retention time reproducibility of better than 0.2 s absolute deviation of retention times to be considered a separate peak. Liners were replaced after each set of 10 injections. Mass spectrometry parameters were as follows: a Leco Pegasus IV mass spectrometer was used with unit mass resolution at 17 spectra/s-1 from 80-500 Da at - 70 eV ionization energy and 1800 V detector voltage with a 230 °C transfer line and a 250 °C ion source.

### Metabolomics Data Analysis

For each sample, 806 metabolites were returned with quantified ions (specific mass-to-charge ratio used to quantify the amount of a target compound) and individual retention indexes (confirming the identity of unknowns by comparing this value to the reference database). These metabolites also have their PubChem and Kyoto Encyclopedia of Genes and Genomes (KEGG) identifiers. Among the 806 metabolites, 223 had known binbase names (Bremer et al., 2023). The actual data are given as peak heights for the quantification ion (m/z) at the specified retention index. To reduce the impact of between-series drift of instrument sensitivity caused by machine maintenance, aging, and tuning parameters, the raw data were normalized to the within-sample sum of all peak heights for all identified metabolites, which was called “mTIC” (analogous to Total Ion Chromatogram). Only identified compounds were included in the mTIC to prevent non-biological contaminants from distorting normalization. Subsequently, it was determined if the mTIC averages were significantly different between treatment groups or cohorts. If these averages were indeed different by p < 0.05, data were normalized to the average mTIC for each group. However, since the mTIC averages between treatment groups were not significantly different, individual metabolites per sample were normalized to the average of all peak heights for all identified metabolites (mTIC_average_). The following equation summarizes the normalization for metabolite *i* of sample *j*

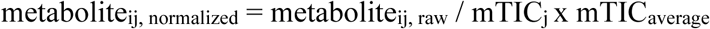

For this study, normalized relative semi-quantified data were used, meaning that the data were peak heights normalized by mTIC_average_. These data were exported into MetaboAnalyst 5.0 (Pang et al., 2021). To conduct a one-factor statistical analysis, we set the low-variance filter to the relative standard deviation (RSD), which is the standard deviation divided by the mean. This is set to filter out 30% of the features as recommended by the software for GC-MS analysis, leaving us with 564 features. Low-variance filtering removes data points that are near constant across all experimental conditions. After filtering, we applied a log_10_ transformation and scaled the data using the autoscaling option (mean-centered and divided by each variable’s standard deviation). We checked that our samples were normally distributed and then proceeded to principal component analysis, performed a fold change analysis, and generated heatmaps of the following variables: unexercised males and females (NEM & NEF), unexercised females /males, PT and TW exercise males /females, and unexercised males and females of DGRP 440 and DGRP 900. We then performed *post hoc* tests to compare individual metabolites for between-treatment differences using Student’s t-test in PRISM.

## Results

### Climbing performance of flies differs across devices and genotypes in males and females

To investigate the effects of exercise devices across multiple genotypes, we exercised males and females from six DGRP-selected genetic lines on the TW and the PT for two hours a day for five days. Our results revealed that device (p<0.0001), genetic line (p<0.0001), and sex (p<0.0001) impacted the climbing performance of the flies **(Table 2)**. We also found there is a highly significant interaction between genetic line and exercise device (p<0.0001). The interaction between genetic line and sex significantly affected climbing performance (p<0.0001), indicating that each genetic line’s performance depends on both sex and device type **(Table 2)**.

**Table 2:** Effects of device, sex, line, and their interactions on climbing performance.

| Source | Nparm | DF | Sum of Squares | F Ratio | Prob > F |
| --- | --- | --- | --- | --- | --- |
| Line | 5 | 5 | 5350.8079 | 113.6472 | <0.0001 |
| Device | 2 | 2 | 356.8518 | 18.9482 | <0.0001 |
| Sex | 1 | 1 | 618.9625 | 65.7315 | <0.0001 |
| Line*Device | 10 | 10 | 1978.2960 | 21.0088 | <0.0001 |
| Device*Sex | 2 | 2 | 83.4954 | 4.4335 | 0.0119 |
| Line*Sex | 5 | 5 | 814.8341 | 17.3065 | <0.0001 |
| Line*Device*Sex | 10 | 10 | 169.6251 | 1.8014 | 0.0550 |
| Total | 35 | 35 | 72693.053 |  |  |

**Table 3:** Effect of device, sex, genotype, and their interactions on the expression of mitochondria and stress genes in DGRP 440, 802, and 805.

| GENE | SEX (S) | GENOTYPE (G) | DEVICE (D) | GxD | GxS | SxD | GxDxS |
| --- | --- | --- | --- | --- | --- | --- | --- |
| *srl* | ns | 0.0012 | ns | ns | ns | ns | 0.0321 |
| *Hsp22* | ns | <0.0001 | ns | 0.0014 | ns | ns | 0.0152 |
| *Gagr* | ns | 0.0001 | 0.0141 | 0.0054 | ns | ns | ns |
| *sid* | ns | <0.0001 | ns | ns | ns | ns | ns |
| *Tspo* | ns | <0.0001 | 0.0309 | 0.0062 | 0.0099 | ns | 0.0306 |
| *Upd3* | <0.0001 | ns | ns | ns | ns | ns | ns |

We found that the natural climbing performance of males was significantly higher than that of females in five genetic lines prior to exercise **(Fig. 2A)** but observed that females of genotype 907 climbed higher before the exercise regimen than males **(Fig. 2B)**. Device effects on climbing performance were consistent between sexes within a given genotype **(Fig. 2C)**. Genotypes 802 and 805 performed significantly better on the TW than the PT in both females and males **(Fig. 2C and 2D)**. The controls on the device (TWC and PTC) showed similar patterns of increased climbing performance in some genotypes **(Fig. 2C and 2D)**, suggesting that the restricted movement (1cm spacing) could still enhance climbing performance in some genotypes. DGRP 440 showed improved climbing performance relative to unexercised controls only after PT training, while DGRP 802 and 805 improved only after TW training (**Fig. 2C and 2D**). Previous findings have established significant effects of genetic variation on post-exercise climbing performance in flies on TW (Watanabe et al., 2020) and PT (Backlund et al., 2025). In our study, genetic variation also strongly drove the differences in climbing performance on PT and TW.

**Figure 2:**
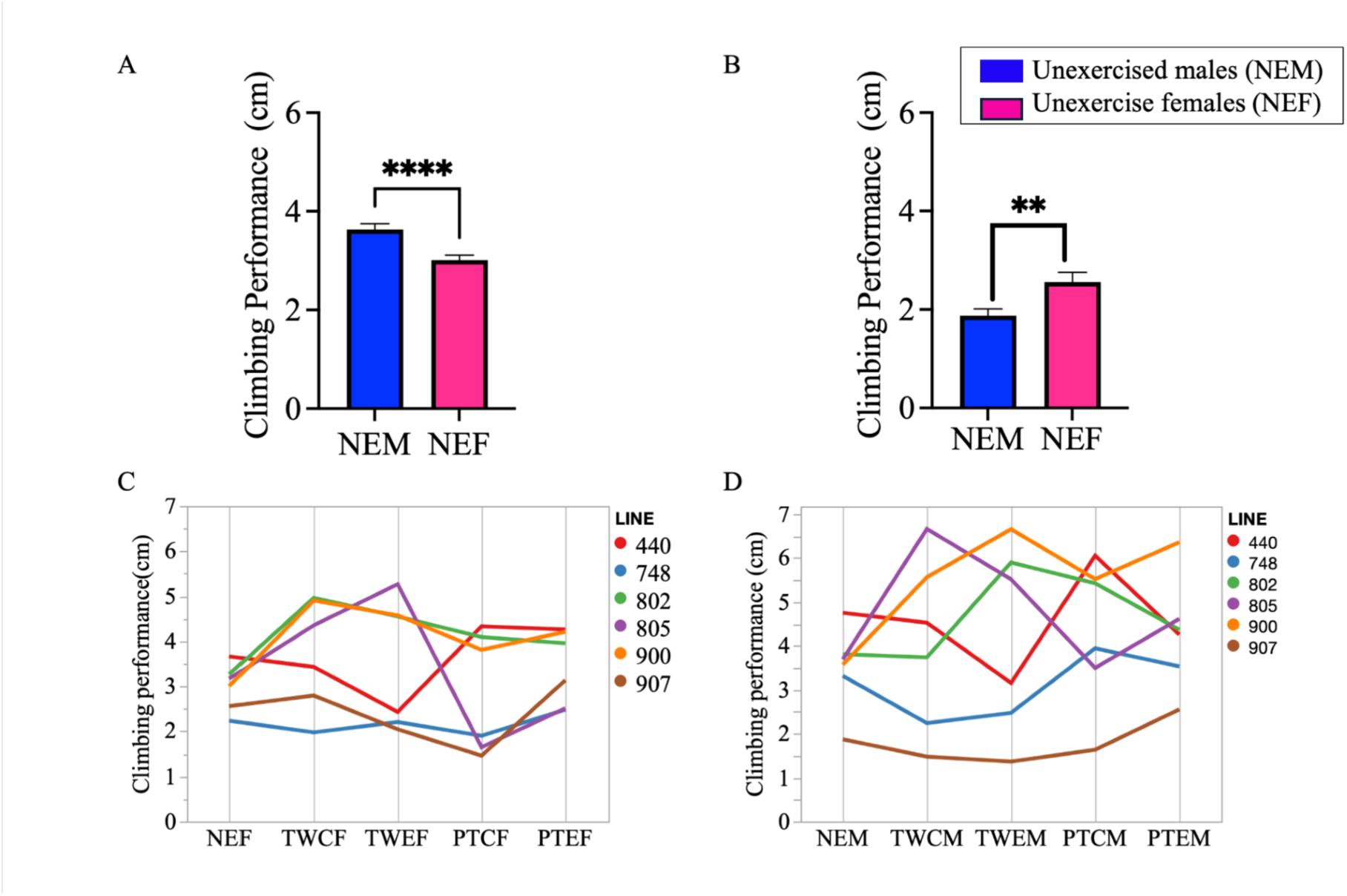
Climbing performance across genotypes, sex, and exercise treatments. **A)** Overall climbing performance across five DGRP lines prior to exercise training. NEM (unexercised males) show naturally high climbing performance in DGRP 440, 748, 802, 805, and 900, with p<0.0001. **B)** Climbing performance of DGRP 907, in which unexercised females (NEF) outperformed unexercised males (NEM; p = 0.001). **C)** Parallel plot showing the mean climbing performance (cm) of females from six genetic lines across device treatments. **D)** Parallel plot showing the mean climbing performance (cm) of males from six genetic lines across device treatments.

### The complex interaction of genotype, device, and sex impacts the expression of stress genes

Following the climbing assay analysis, we determined whether the regulation of the genes involved in mitochondrial function and stress would change depending on the devices on which flies were exercised. We selected DGRP 440 because it showed a higher climbing performance on the PT **(Fig. 2C)**, and we selected DGRP 802 and 805 for the TW because they performed better on the TW **(Fig. 2C)**.

We assessed gene expression after 30 minutes of exercise training. Two genes (*Gagr* and *Tspo*) showed device-dependent differences in expression **(Table 3)**. The main effect of sex was observed only in *Upd3*. Five genes (*srl*, *Hsp22*, *Gagr*, *sid,* and *Tspo*) were differentially expressed across the three genotypes, indicating a significant main effect of genotype. In three genes (*srl*, *Hsp22*, and *Tspo*), we found that the interaction between sex, genotype, and device was highly significant. This means that the expression of these genes was highly determined by the genetic line, the devices flies exercised on, and the sex. Individual gene expression results by the treatment group are shown in **Fig. S1-S6**.

We then conducted a *post hoc* test to identify treatment groups with significantly different expression levels. We observed that following 30 minutes of exercise, females of DGRP 805 that exercised on the TW had a significantly increased expression of *Hsp22* when compared to controls and PT-exercised flies, but there were no changes in DGRP 802 and 440 males and females **(Fig. 3A)**. Additionally, we found that there was a natural variation in *Hsp22* expression across genotypes independent of the treatments. DGRP 440 expressed *Hsp22* more than DGRP 805 and 802, and this pattern was consistent in males and females **(Fig. 3B and 3C)**.

**Figure 3:**
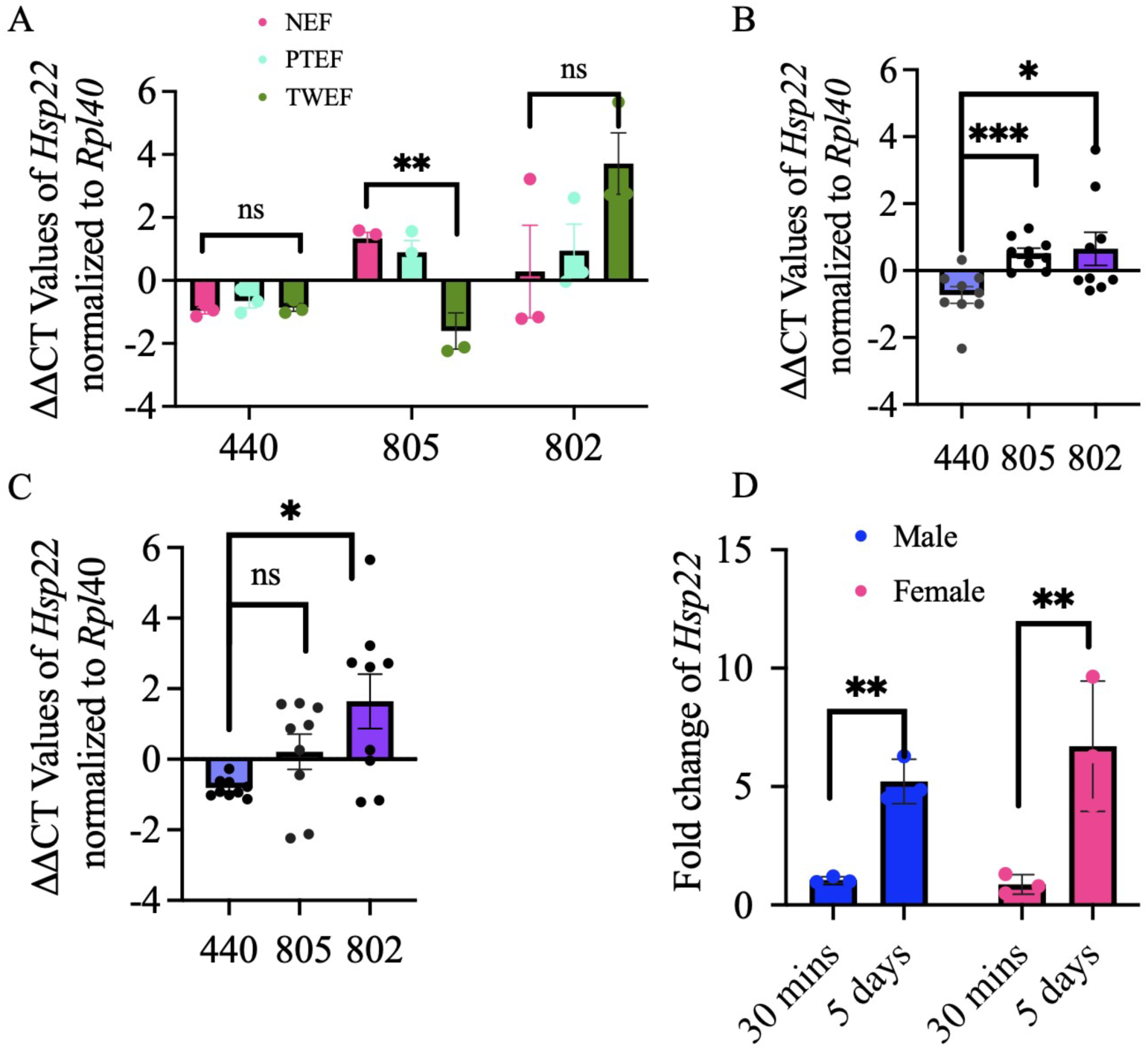
Increased exercise duration upregulated *Heat Shock Protein 22* (*Hsp22*) across both sexes. **A)** Device-specific expression of *Hsp22* in flies that exercised on the TW and PT for 30 minutes, showing increased expression of *Hsp22* in DGRP 805 females that exercised on TW (p = 0.0084). Here, we used three replicates per treatment and n=10 flies per replicate, per genotype. **B)** Expression of *Hsp22* in males across the three genotypes following 30-minute acute exercise, independent of exercise device treatment. *Hsp22* is more expressed in DGRP 440 than in 802 (p = 0.0288) and 805 (p = 0.0008). Here we pooled all treatment groups within each genotype and measured the between-genotype differences. Therefore, we had nine replicates per genotype with n=10 flies per replicate. **C)** Similarly, the expression of *Hsp22* in females revealed between-genotype differential expression of the *Hsp22* gene with increased expression in DGRP 440 (p = 0.0126). We used nine replicates per genotype with n=10 per replicate. **D)** Upregulation of *Hsp22* after five days of continuous exercise training compared to the 30-minute exercise in DGRP 440 males (p = 0.0004) and females (p = 0.0050). The statistical values were determined by conducting an unpaired t-test with Welch’s correction with error bars representing ± SEM (standard error of the mean)

We found that in males, the three genotypes did not differentially express *srl* after exercise on the PT or TW, but females of DGRP 802 showed a significant downregulation of *srl* after exercise on the PT **(Fig. S1)**. *Upd3* was highly expressed in DGRP 440 males that exercised on the TW, relative to females; there were no significant changes in expression in females across all treatments **(Fig. S2)**. Expression of *Gagr was down*regulated in DGRP 805 males and females that exercised on PT and TW compared to controls, while there was no difference in the expression across treatments in DGRP 802 and DGRP 440 **(Fig. S3)**. *Sid* was only detectable in TW males after 30 minutes of exercise **(Fig. S4)**. Also, *Tspo* was upregulated in DGRP 440 males and 805 females but downregulated in 440 females who trained on TW. Lastly, *Hsp22* was expressed at higher levels in DGRP 440 males who exercised on the PT **(Fig. S5)**. The complex interplay among exercise device, sex, and genotype shapes expression patterns of these genes after a 30-minute treatment.

### Downregulation of stress genes after five days of exercise

Next, we conducted a five-day exercise training, during which the flies exercised for two hours each day. We then measured the expression pattern of the genes we previously measured in the 30-minute cohort in DGRP 440. We observed that *Hsp22* expression was significantly upregulated in both devices and sexes after five days of exercise relative to the 30-minute exercise **(Fig. 3D)**.

Conversely, after exercise, the expression of four other genes (*Gagr*, *sid*, *Upd3*, and *vri)* was downregulated in TW exercised males and females **(Fig. 4A and 4B**). *Tspo* expression decreased in males but was not altered in females that exercised on TW **(Fig. 4A and 4B)**. Similarly, five genes (*Gagr*, *sid*, *Upd3*, *drp1,* and *vri*) were downregulated in male flies that exercised for five days on the PT **(Fig. 4C)**. In females who exercised on the PT, only the *Gagr* gene was downregulated after five days of exercise compared to 30 minutes **(Fig. 4D)**. We observed a general pattern of downregulation of these genes in TW-exercised males and females, and PT males. In females on the PT, the expression of other genes except *Hsp22* (upregulated) and *Gagr* (downregulated) was not altered by increasing exercise to five days.

**Figure 4:**
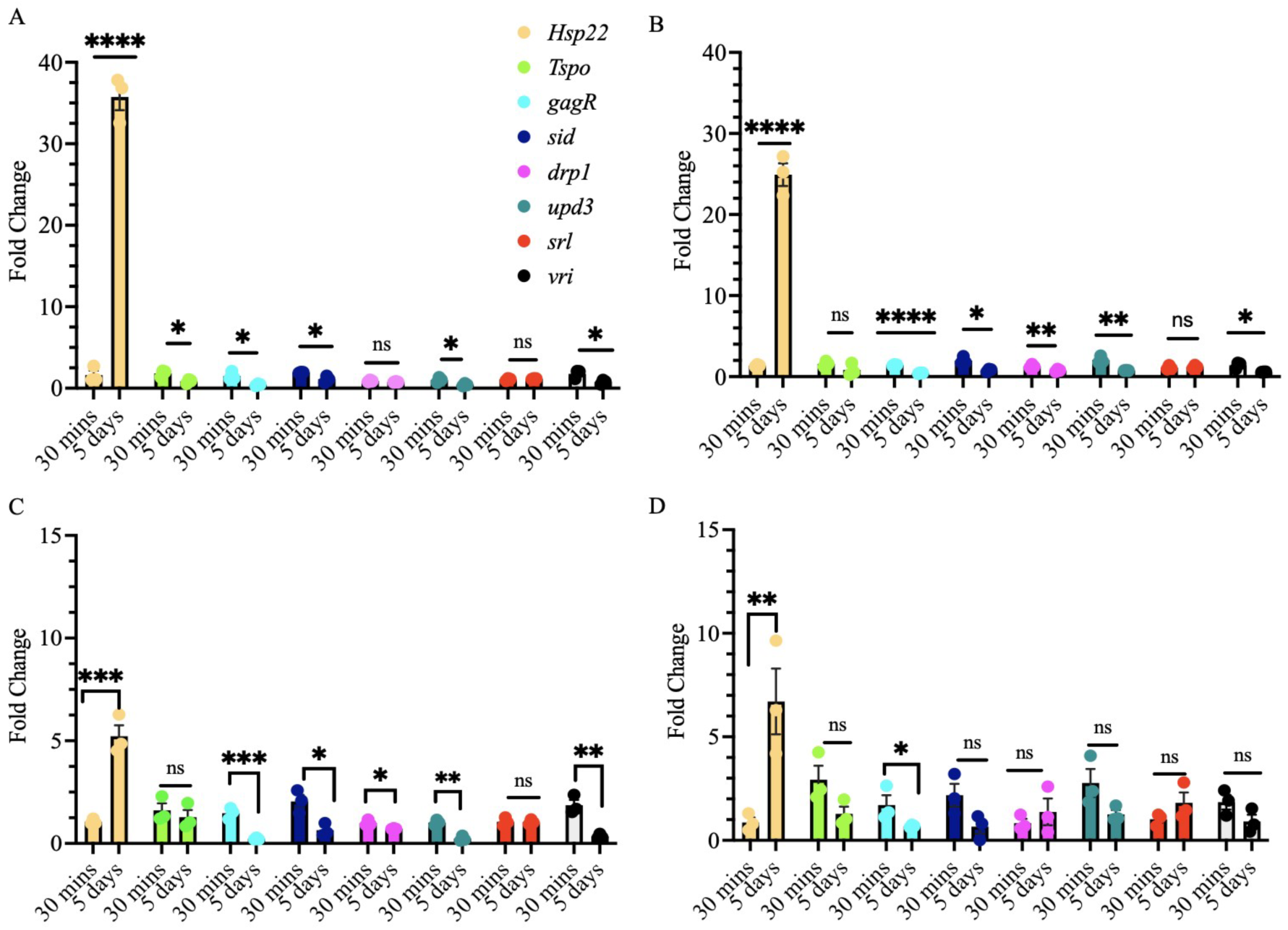
Downregulation of genes associated with stress suggests increased adaptation to the exercise effect after five days of exercise. Change in the expression of eight genes associated with stress, inflammation, and mitochondrial functions after 30 minutes vs. five days of exercise on the TreadWheel for males **(A)** and females **(B)** of DGRP 440 exercised on TW. Change in expression of eight genes linked with stress, inflammation, and mitochondrial function in DGRP 440 males **(C)** and females **(D)** exercised on PT.

### The metabolomes differ for untrained male and female flies of DGRP 440

We further characterized the effects of five days of exercise on PT and TW using untargeted analysis. Using this approach, we integrated the effects of device types, sex, and genotypes on genetic pathways. Given our observation that males’ natural climbing performance was higher than that of females across five genetic lines **(Fig. 2A)**, we examined the metabolome of unexercised males and females to determine whether there is a difference in the abundance of metabolites prior to exercise.

Using principal component analysis, we confirmed that the metabolome of male flies that were not exercised differed significantly from their female counterparts **(Fig. 5A)**. This is consistent with our findings in **Fig. 2A**, which showed that baseline fitness levels of males and females across five genetic lines differed significantly prior to exercise. The heatmap of compounds showed a distinct separation in the concentrations of the first fifty significant metabolite features **(Fig. 5B)**. Asparagine, phenylalanine, glutamine, and guanosine had increased concentrations in females compared to males, while polyunsaturated and monounsaturated fatty acids (myristic acid, stearic acid, linoleic acid, palmitic acid, and linolenic acid), and glycerol had decreased concentrations in females compared to males **(Fig. 5B and 5C)**. The PCA, heat map, and volcano plot buttressed underlying sexual dimorphism in the baseline metabolite profile **(Fig. 5)**.

**Figure 5:**
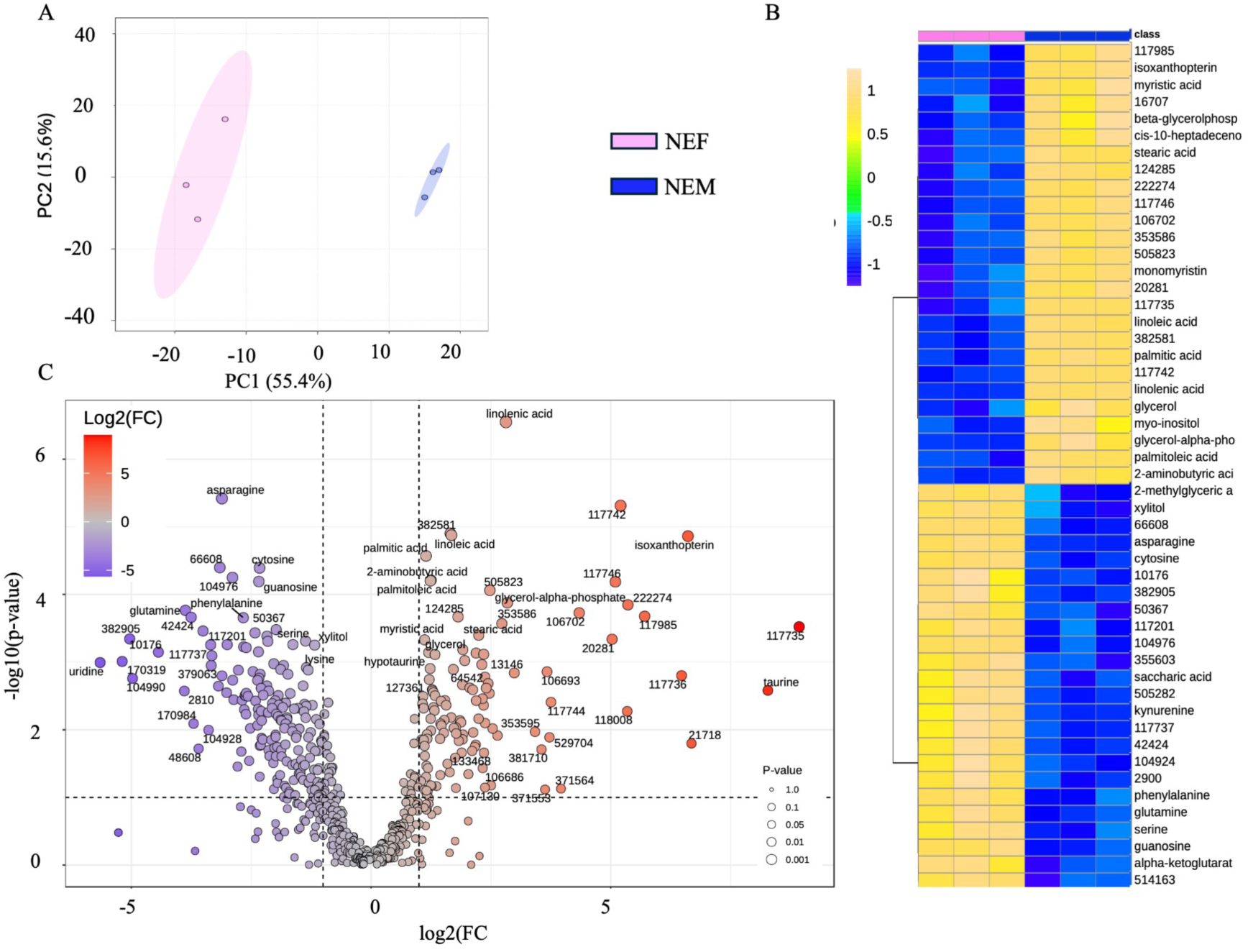
Sexual dimorphism impacts metabolite abundance in untrained flies. **A)** Principal component analysis of the metabolite profiles of DGRP 440 males (NEM) and females (NEF) before exercise training, showing clear clustering by sex. Each point represents a replicate. **B)** Heatmap of fifty metabolites that differ between males (NEM) and females (NEF) of DGRP 440 before exercise. The fold change of the normalized peak heights for each metabolite was determined, and the heatmap of the first fifty significant features was set at p < 0.05. A relative increase is represented by a yellow gradient, while a relative decrease is represented by a blue gradient. **C)** Volcano plot showing the fold change of compounds in males compared to females. The y-axis of the volcano plot shows the statistical significance at –log_10_ P-value, while the sizes of the dots represent the significance levels of the metabolites. The blue color indicates compounds with lower abundance in males but higher in females, while the red color represents compounds with higher fold change in males compared to females.

### The metabolome of flies exercised on the PT is distinctly different from that of unexercised flies

To determine whether the metabolome of flies changes when exercised on the PT and TW, we employed principal component analysis (PCA) to assess whether there is distinct separation between flies that exercised on these devices compared to their controls. Principal component analysis showed distinct clustering of flies that exercised on the PT **(Fig. 6A)**, without a clear separation between the TW-exercised and the unexercised (Fig. 6B). We identified that allantoin, vanillic acid, pipecolic acid, citraconic acid, and aconitic acid, along with other compounds, were elevated in female flies that exercised on the PT (Fig. 6C) but were not detected in males **(Fig. 6D)**. For the TW device, we found that allantoin was also elevated in males and females after exercising on TW, but this was less pronounced than what was observed on the PT **(Fig. 6D, Fig. S6-7)**.

**Figure 6:**
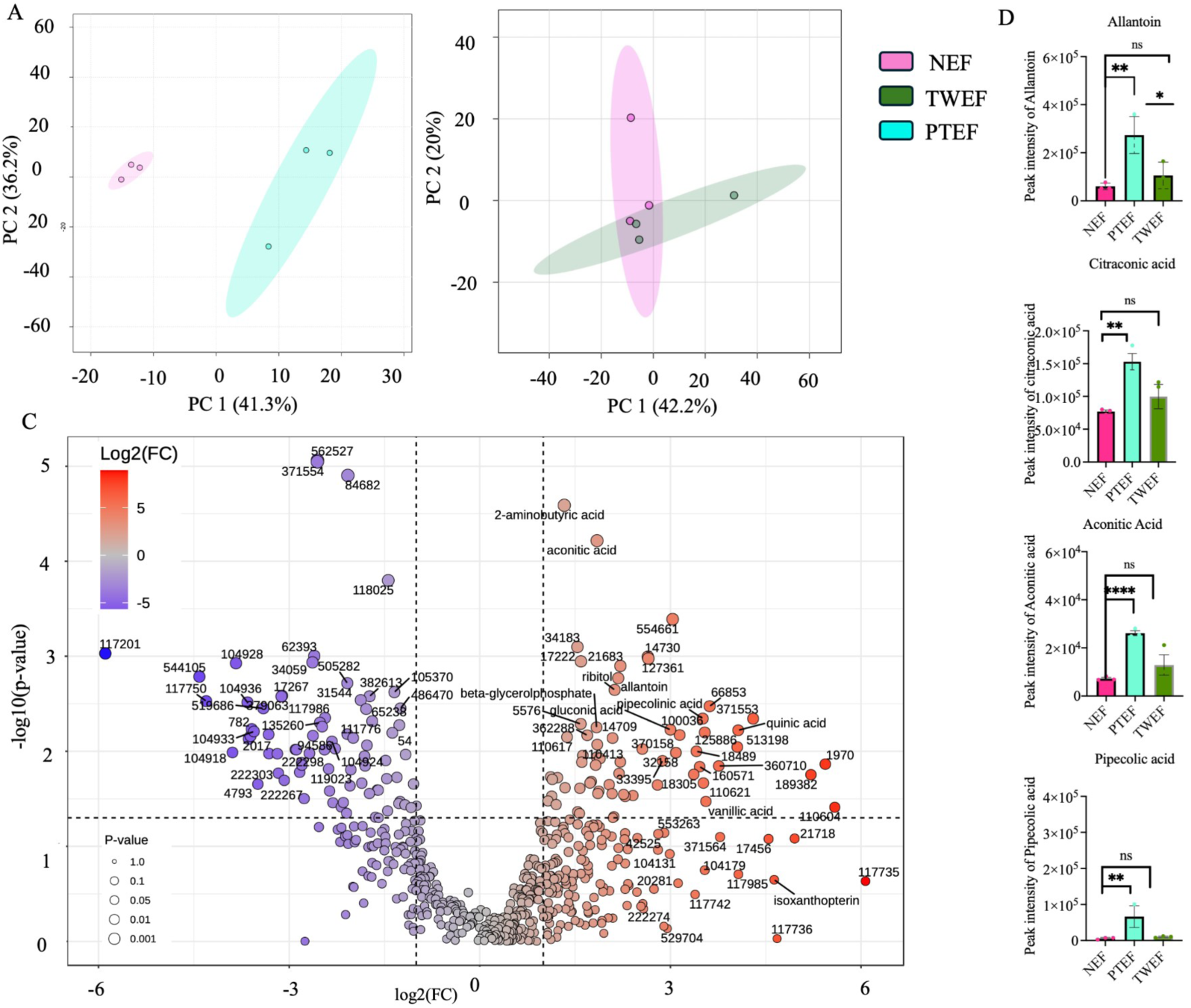
PT induces biomarkers of exercise, oxidative stress, and inflammation in females. **A)** The principal component analysis of NEF and PTEF after five days of continuous exercise on PT shows a distinct clustering pattern. The points represent three replicates per treatment. **B)** The principal component analysis of NEF and TWEF after five days of continuous exercise did not show distinct separation following exercise training on the TW. **C)** Volcano plot comparing the fold change of metabolites of NE and PTEF post-exercise. The y-axis of the volcano plot shows statistical significance as –log10 (p-value), while dot size indicates the significance levels of the metabolites. The blue color shows compounds with lower abundance in PTEF but higher in NEF, while the red color represents compounds with higher concentrations in PTEF compared to NEF. **D)** The peak intensity of some metabolites that are biomarkers of oxidative stress and anti-inflammatory response showed a significant increase in metabolite intensities in PT compared to TW. A post-hoc unpaired t-test was conducted to compare the intensities of these compounds.

To follow up, we compared fold differences in females who exercised on the TW (TWEF) and the PT (PTEF). Most of the metabolites with increased concentration in PTEF relative to NEF were not even detected in TWEF (**Table 4**). The intensities of metabolites like aconitic acid, quinic acid, and pipecolic acid increased on the PT more than on the TW **(Table 5)**. Two metabolites (Gluconic acid and betaglycerophosphate) abundant in PTEF and TWEF showed greater fold change in PT-exercised flies **(Table 5)**.

**Table 4:** Fold change of metabolites linked to oxidative stress and inflammation in PTEF and TWEF compared to NEF.

| Metabolites | Log2FC- PT | Log2FC- TW | Specific roles |
| --- | --- | --- | --- |
| Aconitic acid | 1.855 | Not detected | ATP production, mitochondrial ROS indicator in *Drosophila* (Lushchak et al., 2014), and in mice (Scandroglio et al., 2014) |
| 2 aminobutyric acid | 1.3303 | Not detected | Antioxidant properties in mice (Choe et al., 2021) |
| Quinic acid | 4.0657 | Not detected | Antioxidant properties, anti-inflammatory effects in mice (Lv et al., 2025) and in *Drosophila* (Kim et al., 2025) |
| Vanillic Acid | 3.557 | Not detected | Antioxidant and anti-inflammatory properties in mice (Calixto-Campos et al., 2015) |
| Pipecolinic acid | 3.5078 | Not detected | Antioxidant and anti-inflammatory properties in mice (Zhang et al., 2024). |
| lysine | -2.3135 | Not detected | Oxidative stress indicator and anti-inflammatory properties in mice (Wojtala et al., 2021). |
| Homocysteine | -2.0436 | Not detected | Glutathione synthesis in mice (Yoon et al., 2023). |
| Cysteine | -1.97 | Not detected | Glutathione synthesis in mice (Yoon et al., 2023). |
| Betaglycerophosphate | 1.8375 | 1.6679 | Mitochondrial ROS production in *Drosophila* (Tretter et al., 2017). |

Not detected means these metabolites are not significantly different from the unexercised controls

**Table 5:** Metabolites abundant in untrained males and their relative abundance post-training in TW and PT-trained males vs. females.

| Metabolite | Log2FC Untrained males compared to untrained females | Log2FC PT males compared to PT females | Log2FC TW males compared to TW females |
| --- | --- | --- | --- |
| Linoleic acid | 1.67 | not detected | 1.66 |
| Linolenic acid | 2.82 | 1.73 | 2.50 |
| Myristic acid | 1.11 | not detected | not detected |
| Stearic acid | 2.25 | 1.17 | 1.79 |
| Isoxanthopterin | 6.612 | 2.388 | 5.92 |
| Taurine | 8.27 | 3.86 | 4.19 |
| Asparagine | -3.11 | -1.66 | -3.02 |
| Glutamine | -3.87 | -2.55 | -3.72 |
| Guanosine | -2.34 | -1.23 | -2.04 |
| Cytosine | -2.33 | -1.17 | -1.66 |

Not detected means these metabolites are not significantly different from the unexercised controls

Next, we identified the top metabolites with differential abundance between unexercised males and females (NEM and NEF; **Table 5**). We then compared these to the post-exercise metabolite concentrations in males (PTEM) and females (PTEF; **Table 5**). In NEM, fatty acids like linolenic acid, linoleic acid, and stearic acid were more abundant relative to unexercised females (NEF). Following PT training, linoleic acid was not detected in the PT-exercised males, while the sex difference in other long-chain fatty acids was attenuated in PT-exercised flies. Alongside this, the isoxanthopterin fold change of males relative to females decreased from 6.62 to 2.39 **(Table 5)**, and the log2 fold change of taurine decreased from 8.27 in the untrained to 3.86 in PT-exercised flies. Conversely, the amino acids glutamine and asparagine, along with the nucleosides (guanosine and cytosine), were less abundant in untrained males relative to untrained females **(Table 5)**.

Additionally, to assess the involvement of these metabolites in pathway regulation during exercise, we plotted a network diagram showing the interconnections among metabolic pathways **(Fig. 7A)**. The citrate cycle, purine metabolism, and alanine-aspartate-glutamate metabolism pathways have more metabolites affected by exercise on PT. The color gradient of glutathione metabolism, lysine metabolism, one-carbon pool by folate, cysteine, and methionine metabolism indicated elevated metabolite concentrations in PTEF relative to NEF. We identified four metabolites contributing to glutathione metabolism (L-cysteine, glycine, putrescine, and spermidine), and L-cysteine, a precursor and rate-limiting molecule for glutathione biosynthesis, showed the highest abundance **(Fig. 7B)**.

**Figure 1.7:**
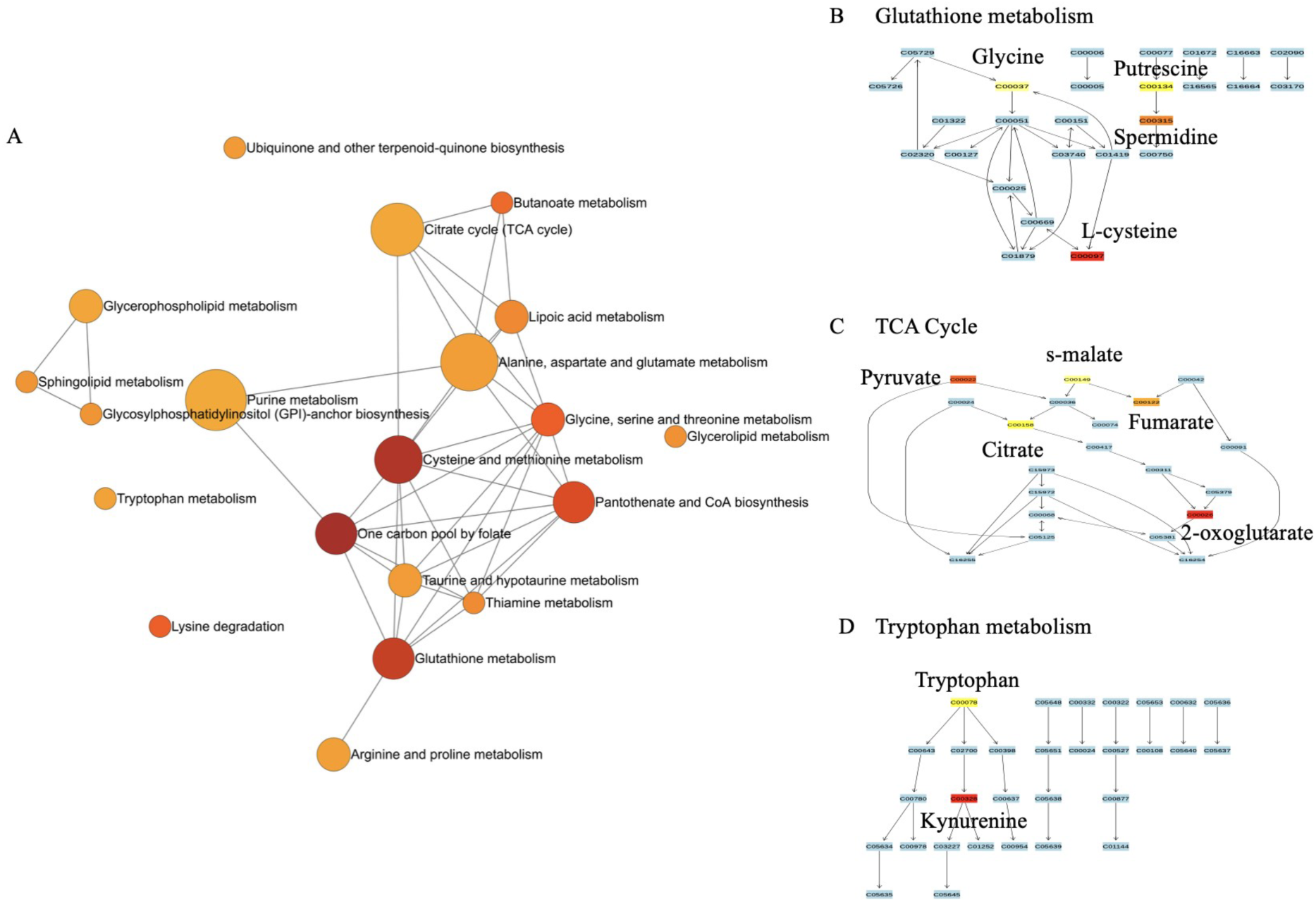
Interactions among pathways for metabolites elevated in female flies exercised on the PT highlight oxidative stress, inflammation, and the TCA cycle. **A)** Network diagram showing the pathway interactions for metabolites that increased in PTEF relative to NEF. Larger nodes indicate that more compounds feed into that pathway. The color of the node indicates the intensity of the metabolites: red indicates nodes whose metabolites in the pathway are highly abundant, while orange and yellow indicate metabolites with lower relative abundance. Schematic diagrams of the KEGG pathway for glutathione **(B)**, TCA cycle **(C)**, and tryptophan metabolism **(D)**, in *Drosophila,* labeled with the names of metabolites identified as elevated in PTEF relative to NEF, colored by amount of elevation (red being greatest).

Five TCA cycle intermediates changed by various degrees in females after PT exercise **(Fig. 7C)**, with five metabolites represented in this pathway. The last pathway highlighted is the tryptophan metabolism pathway **(Fig. 7D)**; its catabolism results in elevated kynurenine levels. Other metabolic pathways are represented in **Table S2**.

Lastly, we determined whether genetic variation contributed to changes in the fly metabolome. First, we compared the metabolome of unexercised flies from DGRP 440 and DGRP 900. The metabolomes of the two genetic lines before exercise were distinctly separated, as shown by principal component analysis for females **(Fig. 8A)** and males **(Fig. S.10)**. The heatmap of the top 25 features shows patterns of metabolite abundance in females, and most of these metabolites are more abundant in 440 NEF than in 900 NEF **(Fig. 8B)**. This suggests that the two genotypes differed in metabolite composition. We observed that, following exercise, the metabolomes of the two genotypes remained distinct **(Fig. 8C)**, but many of the top 25 metabolite features distinguishing the lines after exercise **(Fig. 8D)** differed from those that were differential prior to exercise. One metabolite of note, glucose-6-phosphate, was more abundant in DGRP 900 females than in DGRP 440 females following exercise.

**Figure 8:**
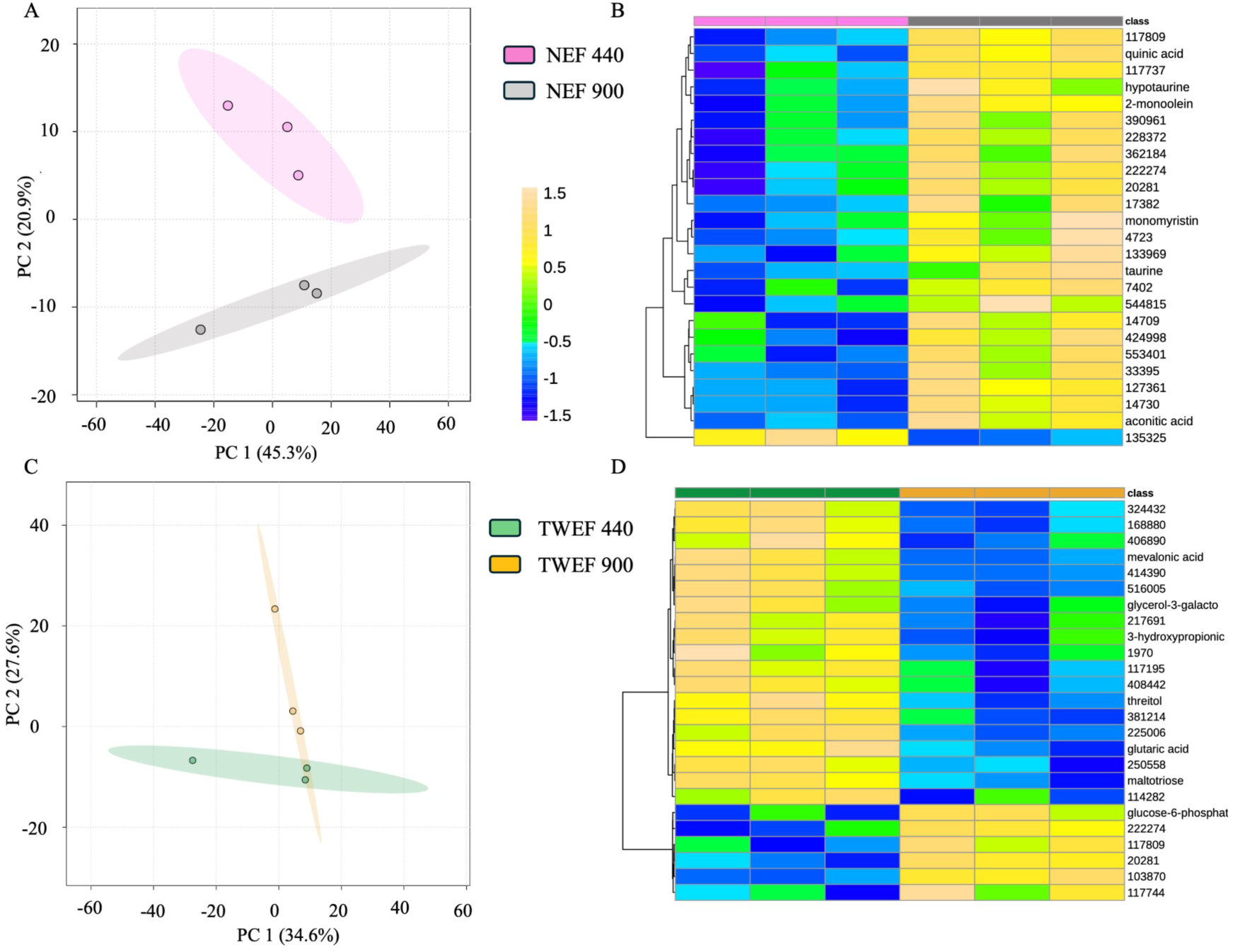
Genetic variation impacts metabolite distribution and response to exercise in females. **A)** Principal component analysis of DGRP 440 and 900 females prior to exercise, showing distinction in the metabolome of flies from different genotypes. **B)** Heatmap of the top 25 metabolite features that differ between DGRP 440 NEF and DGRP 900 NEF. **C)** Principal component analysis of DGRP 440 and 900 females after exercise on TW showing distinct clustering of the samples by genotype. **D)** Heatmap of DGRP 440 TWEF and 900 TWEF highlighting the top 25 distinct metabolic features.

## Discussion

Here, we studied variation associated with exercise in *Drosophila melanogaster* across different devices by comparing climbing performance, gene expression patterns, and metabolome of *Drosophila* that exercised on the Power Tower (PT) and TreadWheel (TW). Our result is consistent with prior studies that have documented extensive genotypic variation in post-exercise climbing performance of flies (Mendez et al., 2016; Watanabe & Riddle, 2021; Backlund et al., 2025).

In our study, some genotypes showed enhanced climbing performance after exercising on a device and had a reverse response on another device. This points to a genuine device-genotype interaction and not a generalized training effect. Device type should be reported alongside genotype and sex when defining exercise responses in *Drosophila*, since the same genotype may respond differently to different devices. Also, this implies that trying different devices could enhance performance for flies that are otherwise not responsive to exercise training on a particular device.

Most observed phenotypic variations are likely underpinned by genetic regulations. In the case of exercise, improved phenotypes (increased running speed, improved cardiac performance, longevity) are associated with the upregulation of several genes in *Drosophila* (Sujkowski et al, 2015). Sujkowski and colleagues found differential expressions of 442 genes in exercise-trained flies that were selectively bred for longevity, and some of these genes activated pathways of glutathione and folate metabolism (Sujkowski et al., 2017). Glutathione pathways regulate stress by scavenging reactive oxygen species to maintain redox balance (Espinosa-Diez et al., 2015). During the acclimation phase of stress, glutathione increases and decreases when stress has caused severe damage (Leeuwenburgh et al., 1995; Izumi et al., 2020). Given that glutathione and JAK/STAT pathways both respond to oxidative stress, and in addition to their overlap in stress regulation, JAK/STAT has been extensively studied in *Drosophila* tissue stress models, we focused on JAK/STAT-associated genes to determine whether exercise elicits a stress response comparable to that elicited by chemical or thermal stressors in *Drosophila* (La Fortezza et al., 2016; Anet et al., 2019; Xin et al., 2020; Sarapultsev et al., 2023; Lv et al., 2024). We studied the expression pattern of *Unpaired 3* (*Upd3*), a cytokine-like protein in *Drosophila,* and a key ligand in the JAK/STAT pathway. We also studied the *Gagr* gene, a retroviral element that especially induces the JAK/STAT pathway to regulate stress (Makhnovskii et al., 2020; Harrison et al., 2021; Balakireva et al.,2024).

In DGRP 802 males, *Upd3* expression was modestly elevated after TW exercise, while *Gagr* was downregulated in DGRP 805 males and females on both devices. This genotype-specific response mirrors what was observed in climbing performance, where similar genotype-device interaction shaped the outcome. Contrary to our expectations, the *Gagr* gene was downregulated, *Upd3* was only slightly upregulated during exercise, and there were mostly no significant changes in the expression of other treatments; however, these two genes were highly upregulated by chemical stressors and infection in *Drosophila* (Nefedova et al.,2022; Gigin et al., 2023).

The weak expression of both genes after 30 minutes may reflect the limitations of acute exercise on gene expression, consistent with evidence that a single exercise bout does not sufficiently alter the expression of some genes in *Drosophila* (Kim et al., 2018). We increased exercise duration to five days while keeping the two-hour regimen constant each day. This time, we narrowed our investigation to one genotype (DGRP 440). The *Gagr* gene was further downregulated across devices and sexes. *Upd3* was also downregulated in exercised flies (males and females) on the TW and in males that exercised on the PT. The progressive downregulation of *Gagr*, *Upd3*, and *sid* following five days of exercise is consistent with transcriptional adaptation, which means that as flies acclimate to repeated exercise bouts, the acute stress signal appears to diminish. A recent study in mice also found that continuous endurance training is associated with pronounced downregulation of stress genes, which is a result of exercise-induced adaptation to stress (De Smalen et al., 2025). Human studies also found a transcriptomic shift from stress, damage, and inflammation to energy metabolism and antioxidant pathways following continuous training compared with acute exercise, which triggers strong stress-related expression (Damas et al., 2018; Bieter et al., 2024).

*Hsp22* expression tells a different story. Unlike *Gagr*, *Upd3,* and *sid*, *Hsp22* was upregulated after five days of exercise in both sexes and across both devices, and the increase was greater than observed after only one bout of exercise for 30 minutes. By measuring *Hsp22* expression, we provided evidence that longer exercise duration increases stress levels. The dual role of *Hsp22,* which could be protective at moderate levels (Morrow et al., 2016) but indicative of cellular stress under chronic overexpression (Morin et al., 2019), presents an argument on whether five days of exercise approaches a beneficial or damaging threshold. Our data reveal the upregulation of *Hsp22* without a corresponding decline in fitness. This suggests that increased stress levels, as depicted by *Hsp22* expression, further support the argument that exercise is protective against stress. Taken with the downregulation of *Gagr*, *Upd3* and *sid*, the *Hsp22* data suggest that exercise simultaneously suppresses acute stress signaling and activates a heat-shock mediated protective response (two processes that are not mutually exclusive).

Gene expression alone was insufficient to differentiate the metabolic consequences of PT vs. TW training, and the metabolome provided that resolution, as we found that exercising on the PT induced distinct changes in the metabolome of male and female flies, more than observed in the TW. We identified several pathways linked to antioxidation and anti-inflammation. Some of these pathways included glutathione metabolism, TCA cycle, lysine degradation, tryptophan metabolism, and energy metabolism (Wu et al., 2004; Martínez-Reyes and Chandel, 2020; Seo and Kwon, 2023). We also identified metabolites directly and indirectly linked to these pathways. These metabolites included allantoin, aconitic acid, citraconic acid, pipecolic acid, quinic acid, and ribitol, among others, detailed in our results section **(Table 4-5)**. In the glutathione pathway, L-cysteine was highly enriched in PT females after five days of exercise compared to unexercised controls. L-cysteine is the building block for glutathione and the rate-limiting molecule of glutathione biosynthesis (Paul et al., 2018). In human studies, increased levels of L-cysteine enhance antioxidant production thereby protecting against oxidative stress (Wu et al., 2004; Averill-Bates et al., 2023; Tan et al., 2023). Our data reflect the exercise effect on the bidirectional relationship between stress and inflammation, as excess reactive oxygen species can increase pro-inflammatory cytokines, then increased antioxidants would further enhance anti-inflammation (Biswas et al., 2006). The elevated levels of citraconic acid and pipecolic acid imply that pathways of antioxidation and anti-inflammation are triggered, especially since citraconic acid is a strong activator of NRF2 pathway of anti-inflammation (Chen et al., 2022; Cheng et al., 2026). In fact, increased citraconic acid and pipecolic have been linked to reduced inflammation and stress in mice (Li et al., 2020; Chen et al., 2022; Cheng et al., 2026).

The extent of these metabolic shifts varied by device, sex, and genotype. We found that the metabolomes of unexercised male and female controls were distinct, and after exercise, the metabolome of flies that exercised on PT (males and females) clearly separated compared to those that exercised on the TW. Most of these metabolites identified above were more abundant in PT females. There was also a genotype-specific difference in the metabolome of untrained flies from DGRP 440 and 900. Exercising further altered metabolite abundance (glucose-6-phosphate, threitol, mevalonic acid) in a genotype-dependent manner. The genotype-dependent metabolome differences observed here are consistent with prior work showing that untargeted metabolomics captures genotype-by-environment interactions in *Drosophila* (Rattray et al., 2018; Reed et al., 2014; Jacob et al., 2019) and extend that framework to exercise as the environmental variable. In this study, it became apparent that factors beyond individual effects are important; the interactions of exercise with other factors are crucial in determining the effectiveness of exercise.

## Conclusion

This study identified the relevance of device type on exercise outcomes. The extent of the effect was dependent on other factors (genotype, sex, and exercise duration). Exercising induced a stress response, which was blunted as flies adapted to exercise over five days. Downregulation of stress genes, in addition to an increase in antioxidant compounds, provided some clarity into the process of exercise-induced adaptation during stress. The *Drosophila* exercise metabolome is a valuable tool for visualizing and interpreting exercise outcomes, as it provides a snapshot of the distinct clustering by sex and device. Our study correlated fitness traits with metabolomic outputs. Most of the metabolites identified as changing with exercise in our study have also been highlighted in human exercise metabolome studies (Starnes et al., 2017; Parstorfer et al., 2025). Our study further confirms that *Drosophila* can be a relevant model organism for exercise research. A limitation of this study is the lack of a direct comparison of the metabolome of the acute and chronic exercise groups. We also did not account for the effects of exercise on feeding preference and its impact on metabolite concentrations. Riddle and colleagues’ findings suggested that 5-day exercise interacts with other factors (sex and genotypes) to mildly impact food consumption and inferred that observed changes in body composition and weight post-exercise cannot be attributed to food consumption alone (Johnson et al., 2024)

Through this study we identified the complex interaction of device, sex, and genotype on gene expression and metabolism and the effect of these interactions on exercise outcomes. Our study revealed that stress can be induced by high intensity exercise over multiple days. *Drosophila* serves as a model organism to better understand human responses, allowing us to draw meaningful conclusions that provide insight into the beneficial outcomes of exercise.

## Acknowledgements

We appreciate the contributions of Michelle Tan (a former undergraduate of the Reed Lab) for helping in data collection during the preliminary phase of this project. We would also like to thank all the Reed Lab members for their contributions and feedback.

## Authors Contribution

Conceptualization: LKR, TRM

Experiment: TRM, AB, JB, MC, ST, AM, AK, MM, CH, GB, JM,

Writing: TRM

Manuscript review and editing: LKR, AB, JB, MC, ST, AM, AK, MM, CH, GB, JM

## Supplemental file

**Figure S1:**
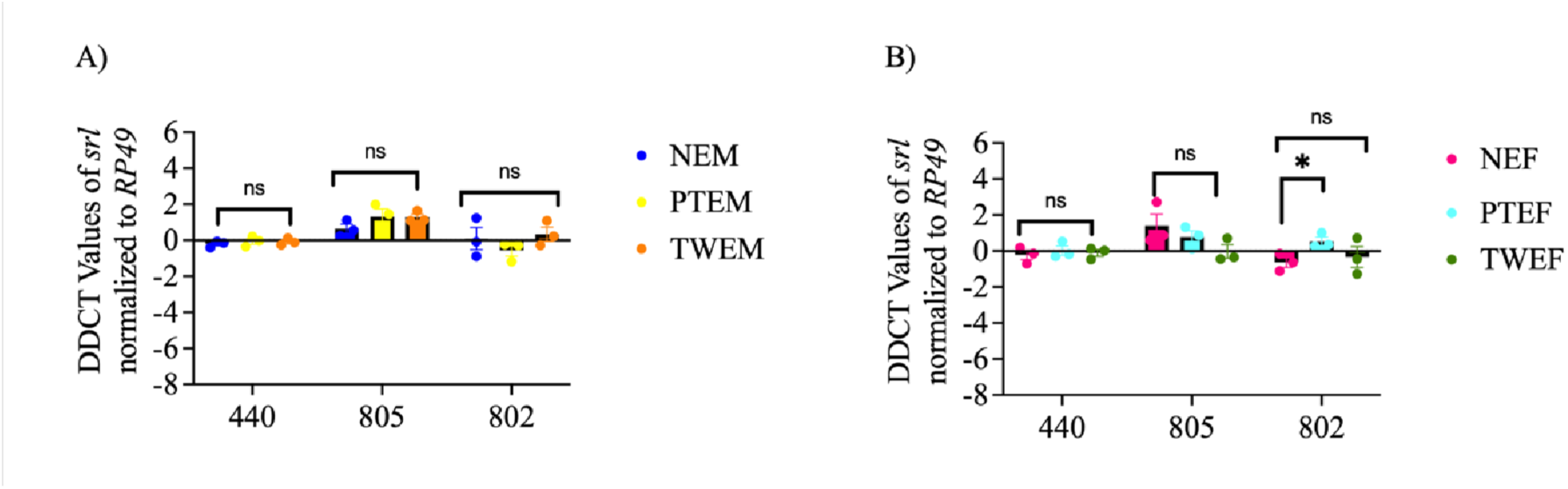
Expression of *spargel* in males and females after acute exercise. **A)** Expression of *spargel* (*srl)* in males after 30 minutes of exercise on TW (TWEM) and PT (PTEM) across three genotypes. **B)** Expression of *spargel* (*srl*) in females following 30 minutes of exercise on TW (TWEF) and PT (PTEF) across three genotypes.

**Figure S2:**
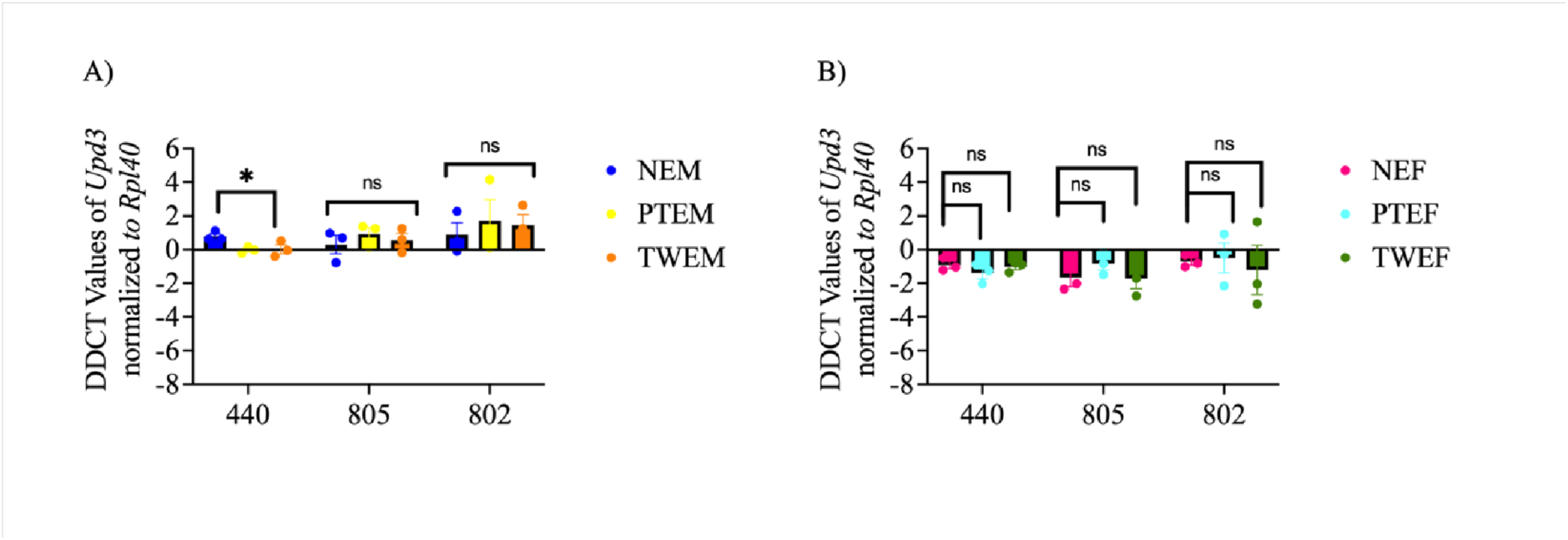
Expression of *Upd3* in males and females after acute exercise. **A**) Expression of *Unpaired 3 protein* (*Upd3*) in males after 30 minutes of exercise on TW (TWEM) and PT (PTEM) across three genotypes. **B)** Expression of *Unpaired 3 protein* (*Upd3*) in females after 30 minutes of exercise on TW (TWEF) and PT (PTEF) across three genotypes.

**Figure S3:**
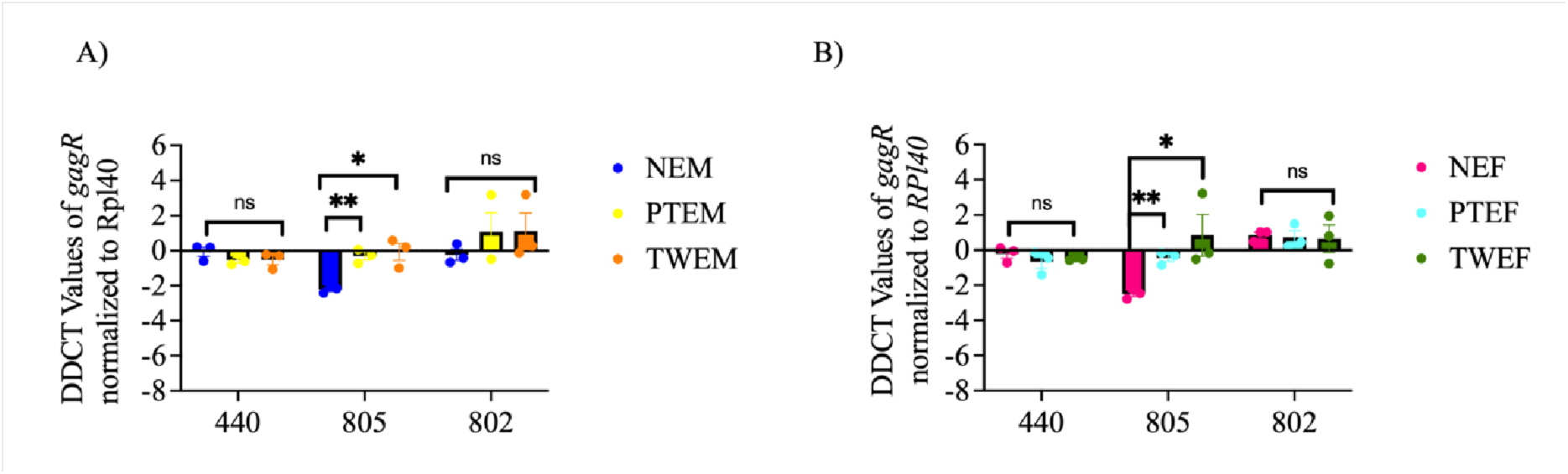
Expression of *gag-related gene* (*Gagr*) in males and females after acute exercise. **A**) Expression of *gag-related gene* (*Gagr)* in males after 30 minutes of exercise on TW (TWEM) and PT (PTEM) across three genotypes. **B)** Expression of *gag-related gene* (*Gagr)* in females after 30 minutes of exercise on TW (TWEF) and PT (PTEF) across three genotypes.

**Figure S4:**
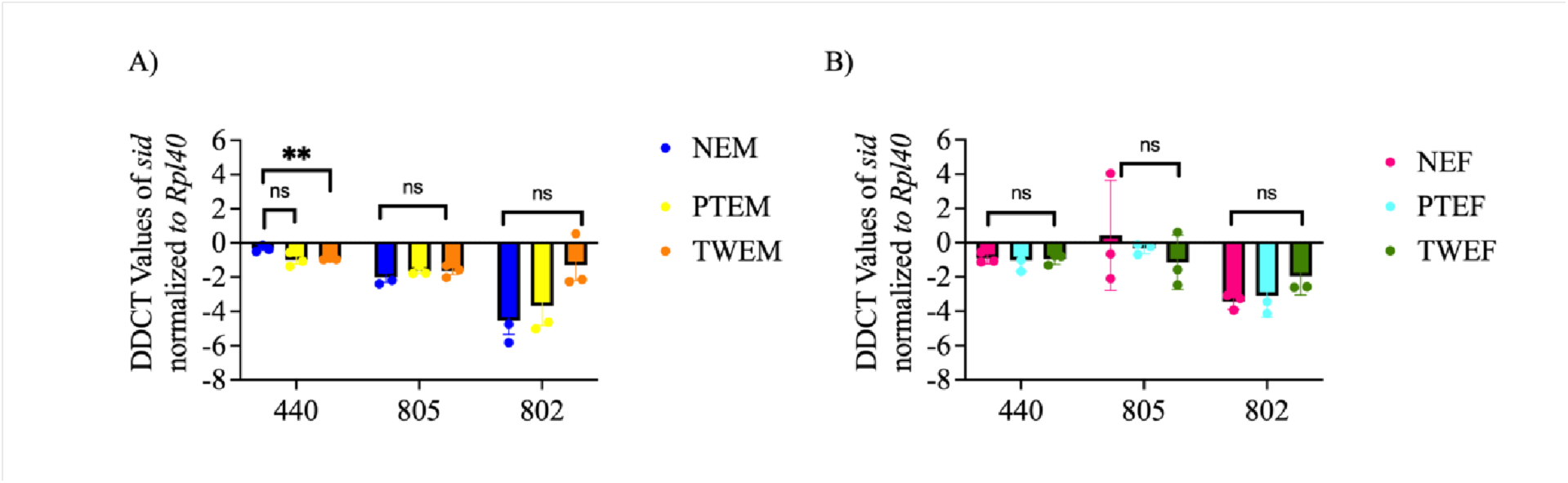
Expression of *Stress-induced nuclease* (*sid*) in males and females after acute exercise. **A**) Expression of *Stress-induced nuclease* (*sid)* in males after 30 minutes of exercise on TW (TWEM) and PT (PTEM) across three genotypes. **B)** Expression of *Stress-induced nuclease* (*sid)* in females after 30 minutes of exercise on TW (TWEF) and PT (PTEF) across three genotypes.

**Figure S5:**
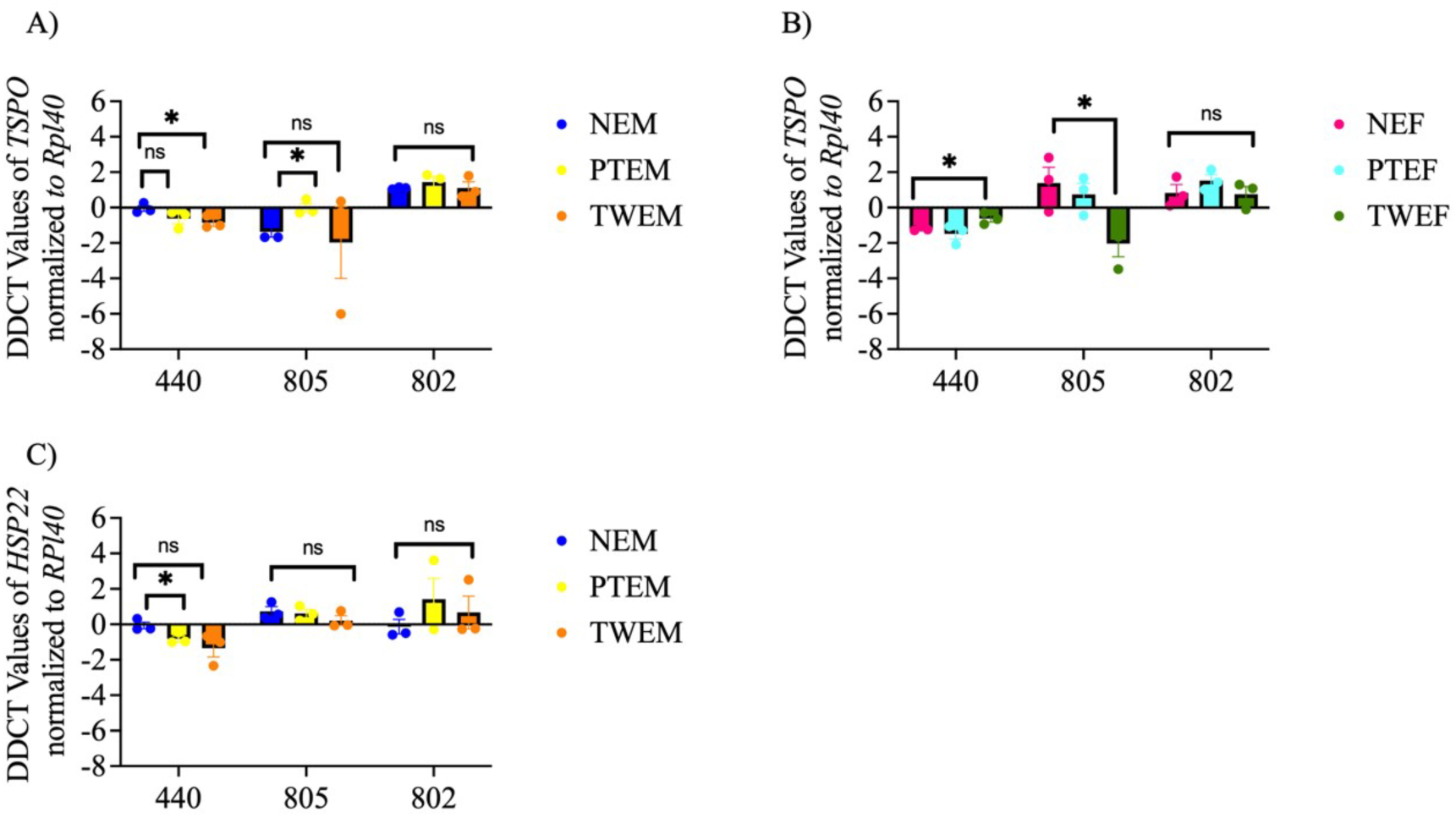
Expression of *Translocator Protein O* (*TSPO*) in males and females after acute exercise. **A**) Expression of *Translocator Protein O (TSPO)* in males after 30 minutes of exercise on TW (TWEM) and PT (PTEM) across three genotypes. **B)** Expression of *Translocator Protein O* (*TSPO)* in females after 30 minutes of exercise on TW (TWEF) and PT (PTEF) across three genotypes. **C)** Expression of *Heat shock protein 22* (*HSP22*) in males after 30 minutes of exercise on TW (TWEM) and PT (PTEM) across three genotypes.

**Figure S6:**
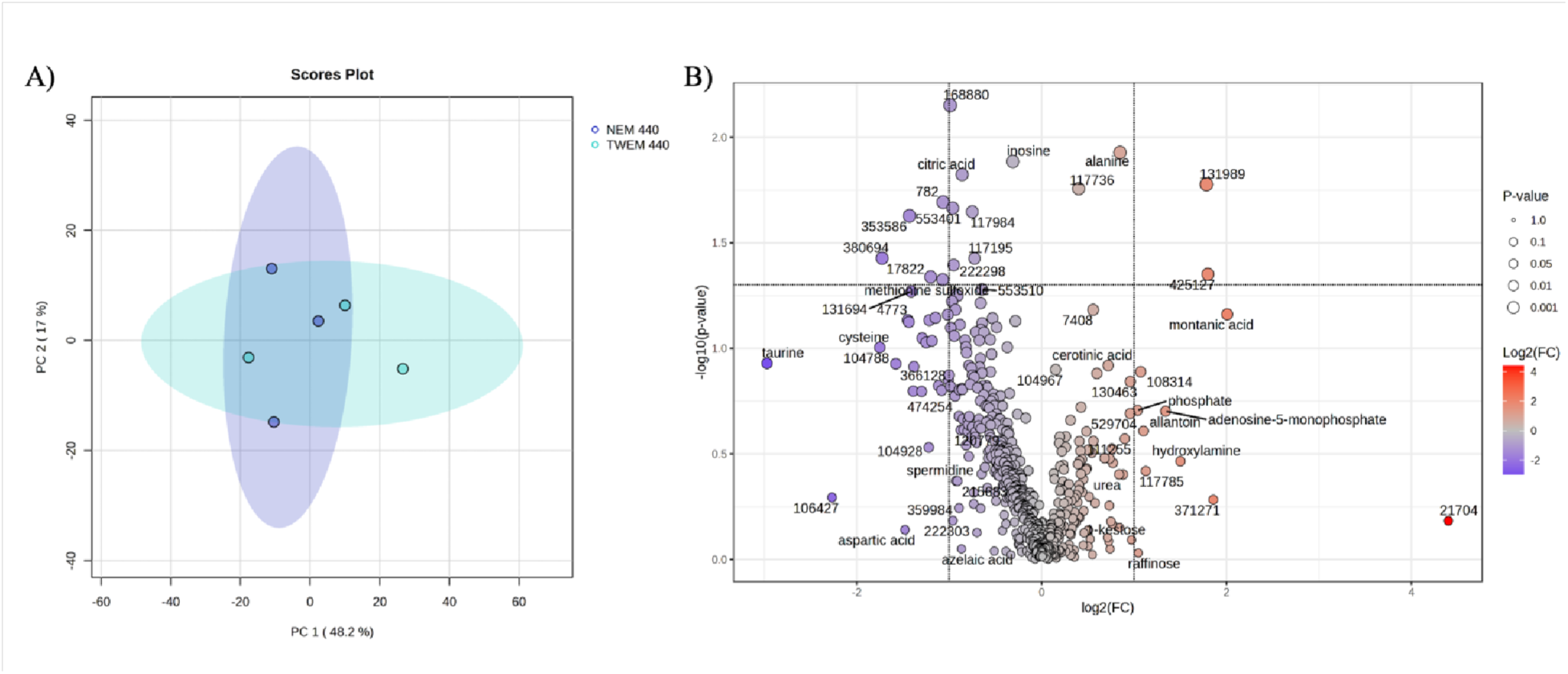
Principal component analysis and differential concentrations of the metabolite profiles of DGRP 440 males on the TreadWheel. **A**) Principal component analysis of the metabolite profiles of DGRP 440 males (NEM) and TreadWheel males (TWEM) after five days of exercise, showing no distinct clustering. **B)** Volcano plot showing the log 2-fold change of compounds in unexercised males compared to TreadWheel males (TWEM). The y-axis of the volcano plot shows the statistical significance at –log10p, while the dot sizes also represent the significance levels of the metabolites. The blue color shows compounds with lower abundance in NEM but higher in TWEM, while the red color represents compounds with higher fold change in TWEM compared to NEM.

**Figure S7:**
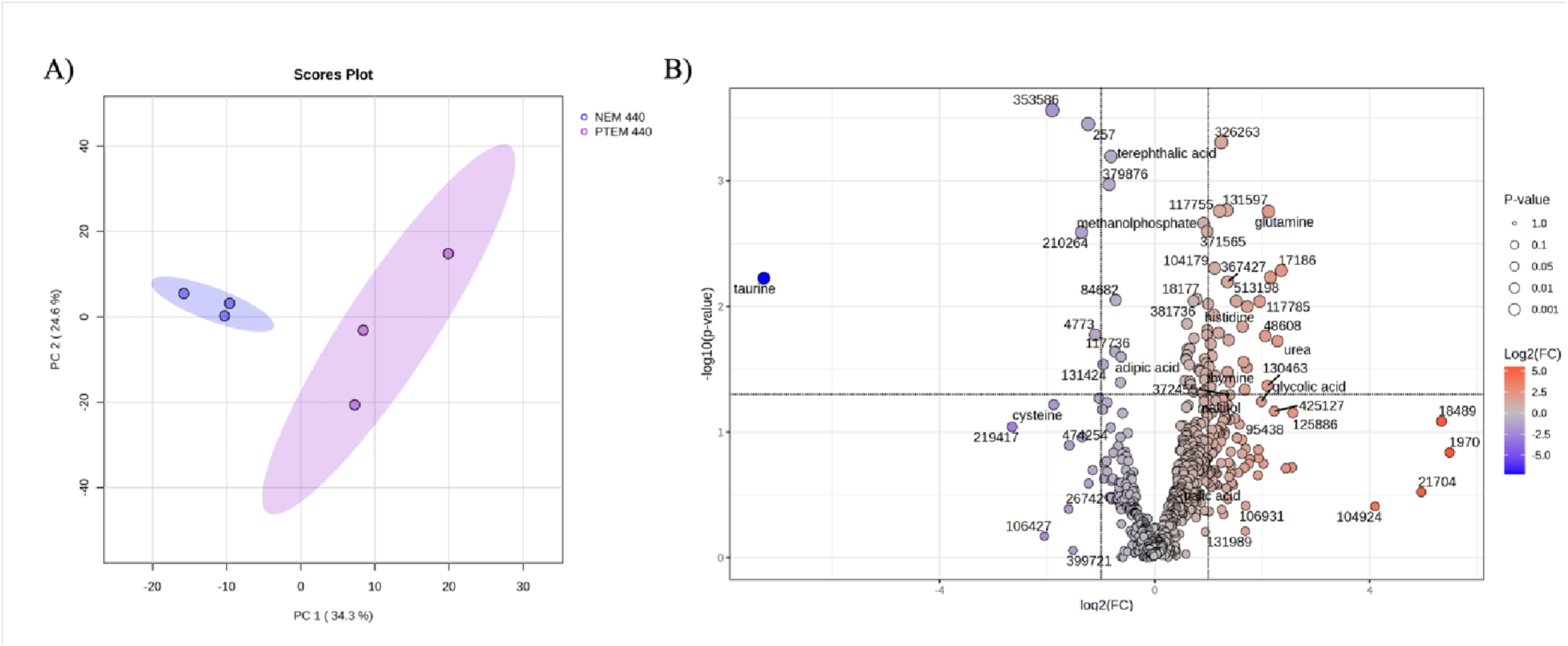
Principal component analysis and differential concentrations of the metabolite profiles of DGRP 440 males on the Power Tower. **A**) Principal component analysis of the metabolite profiles of DGRP 440 males (NEM) and Power Tower males (PTEM) after 5-day exercise, showing distinct clustering. **B)** Volcano plot showing the log 2-fold change of compounds in unexercised males compared to Power Tower males (PTEM). The y-axis of the volcano plot shows the statistical significance at-log10p, while the dot sizes also represent the significance levels of the metabolites. The blue color indicates compounds with lower abundance in PTEM but higher abundance in NEM, while the red color indicates compounds with higher fold change in PTEM than in NEM.

**Figure S8:**
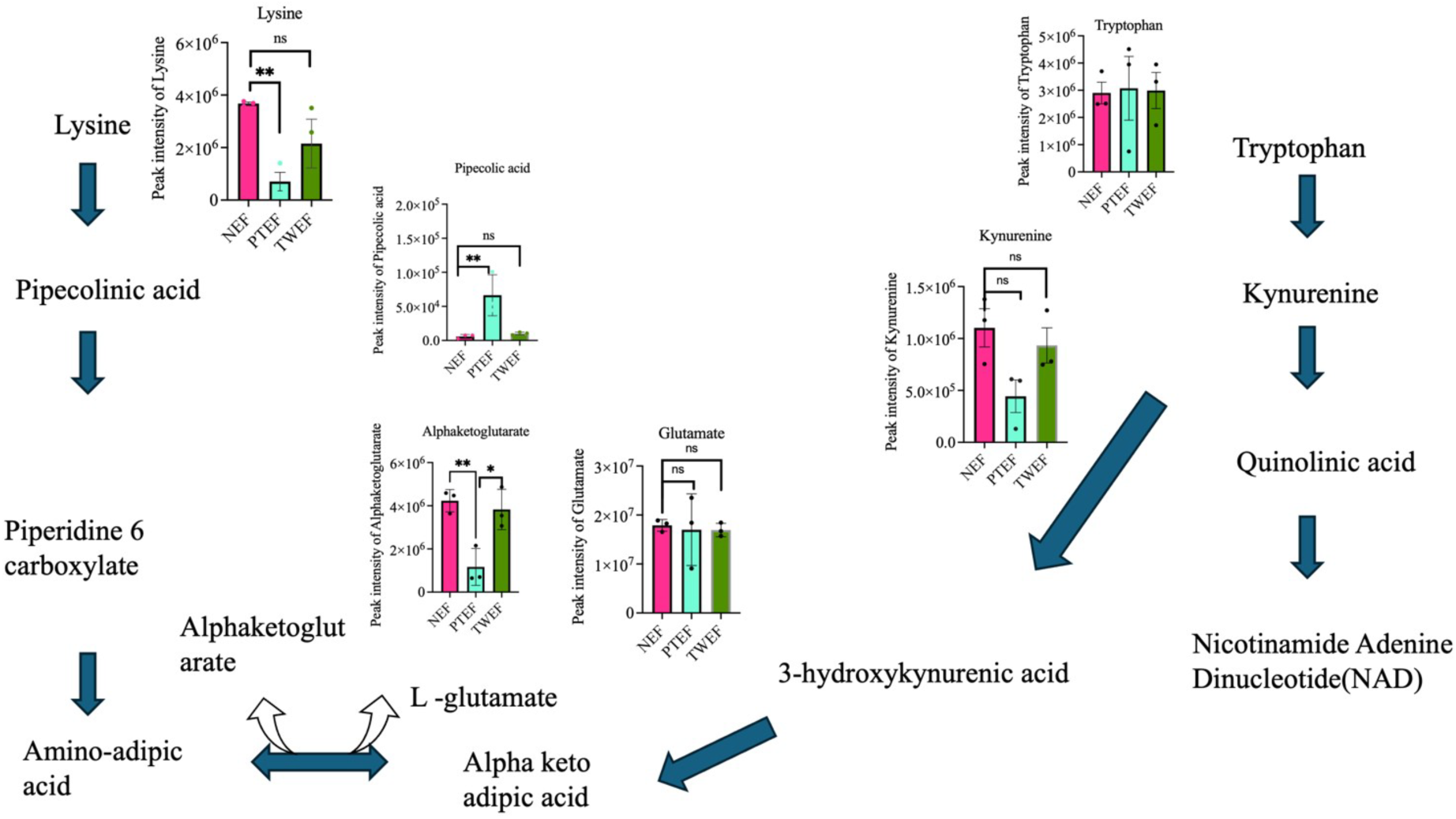
Lysine degradation and tryptophan metabolism pathway co-interaction in females on the Power Tower and TreadWheel. Lysine degradation and tryptophan metabolism pathway co-interaction, highlighting the abundance of certain metabolites in females after five days of exercise on Power Tower (PTEF) and TreadWheel (TWEF).

**Figure S9:**
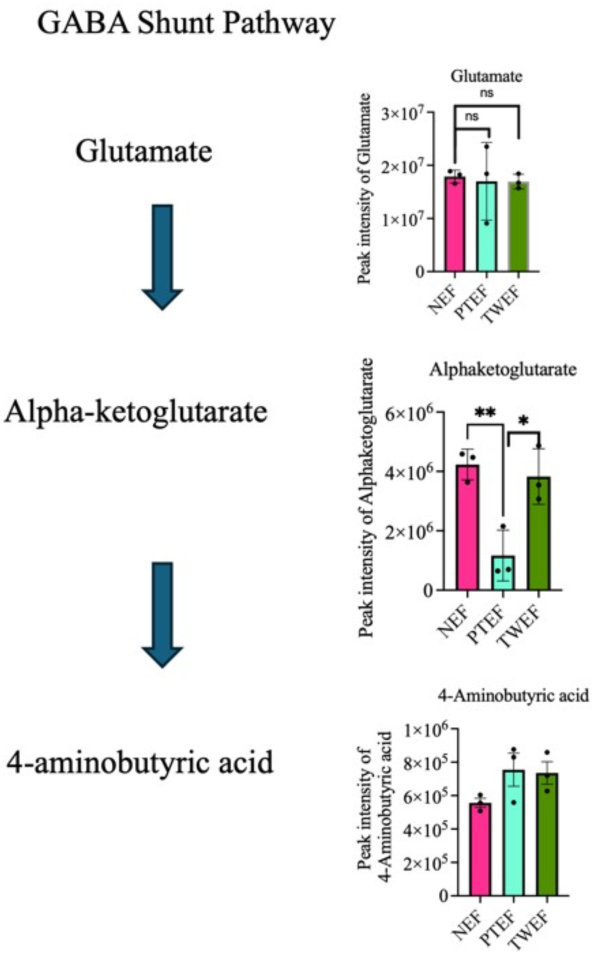
**GABA shunt pathway highlighting some metabolites’ abundance in females after 5-day exercise training.**GABA shunt pathway highlighting some metabolites’ abundance in females after 5-day exercise training on A) Power Tower (PTEF) and B) TreadWheel (TWEF).

**Figure S10:**
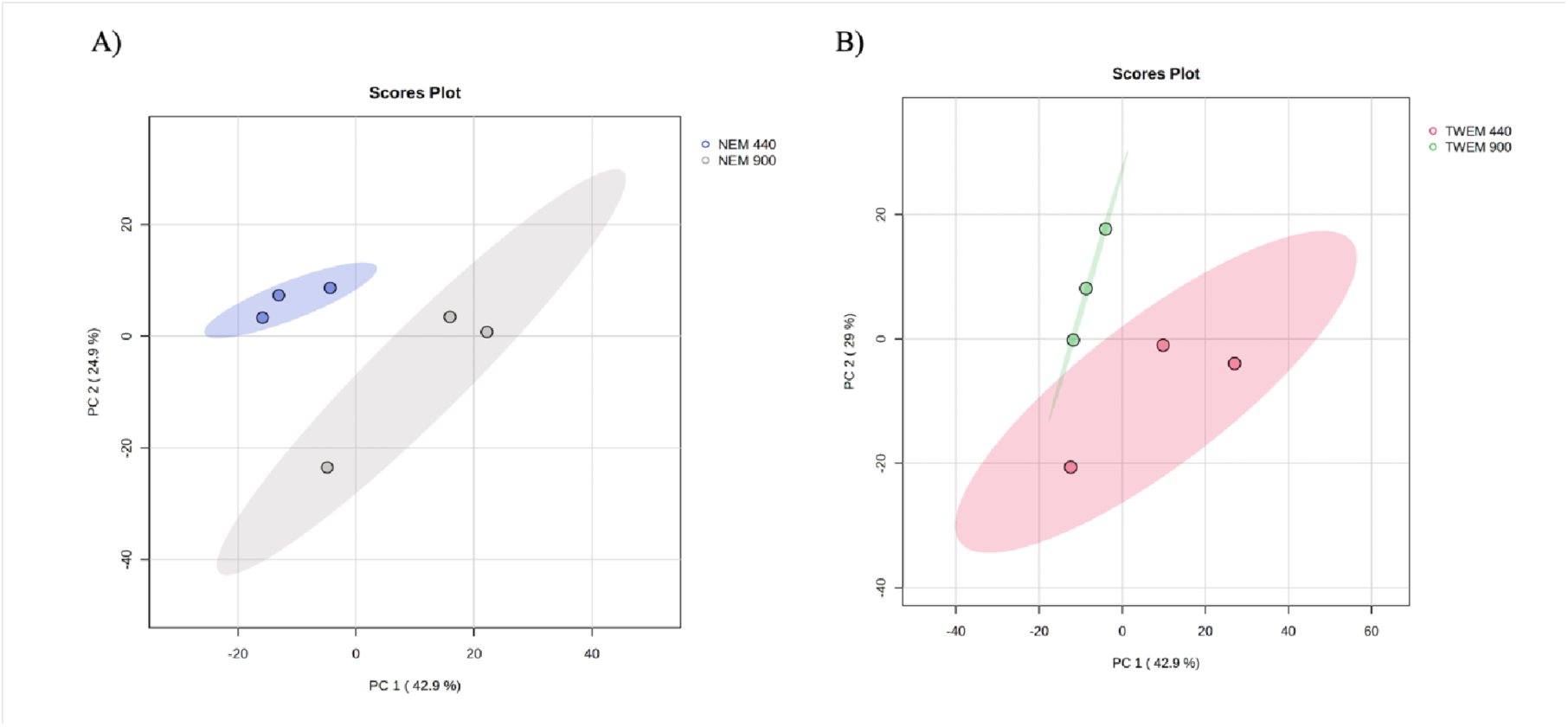
Principal component analysis of the metabolite profiles showing clustering by genotypes. **A)** unexercised males of DGRP 440 and DGRP 900 **B)** TreadWheel-exercised males of DGRP 440 and DGRP 900 males

---

**Table 1.S1:**
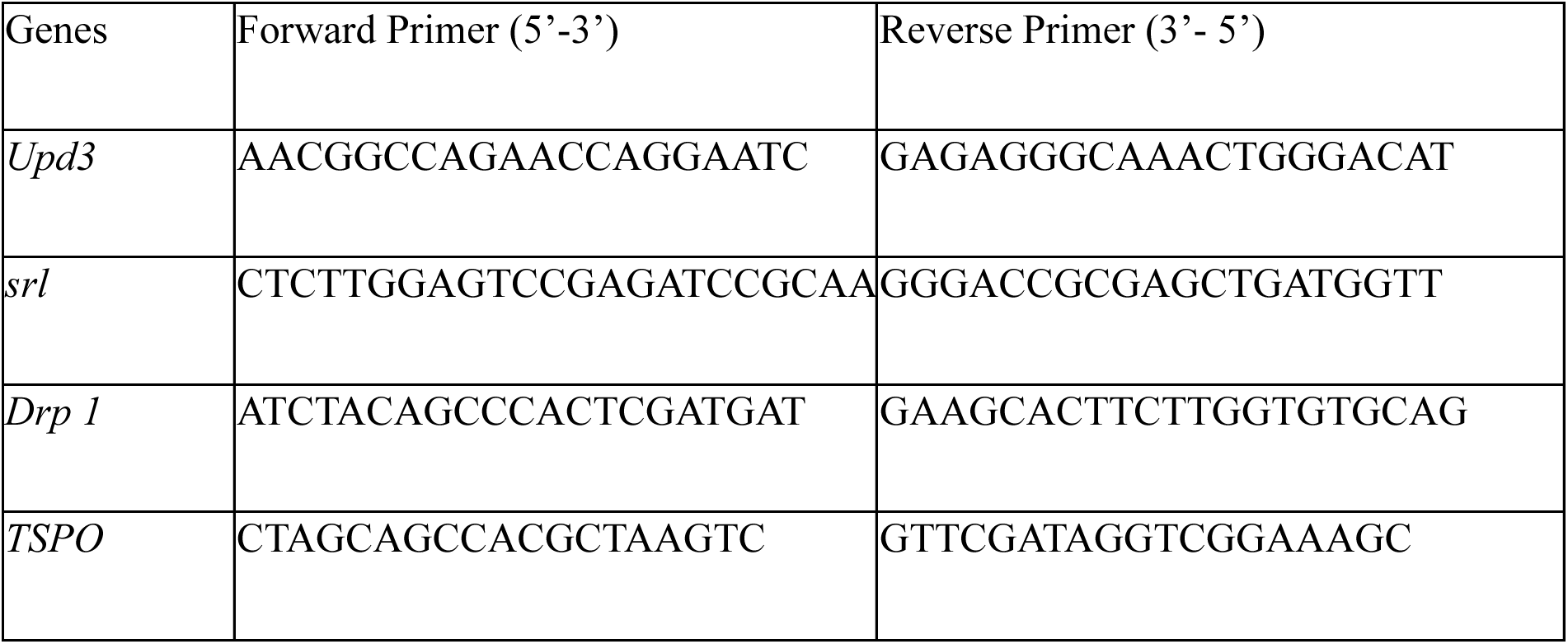
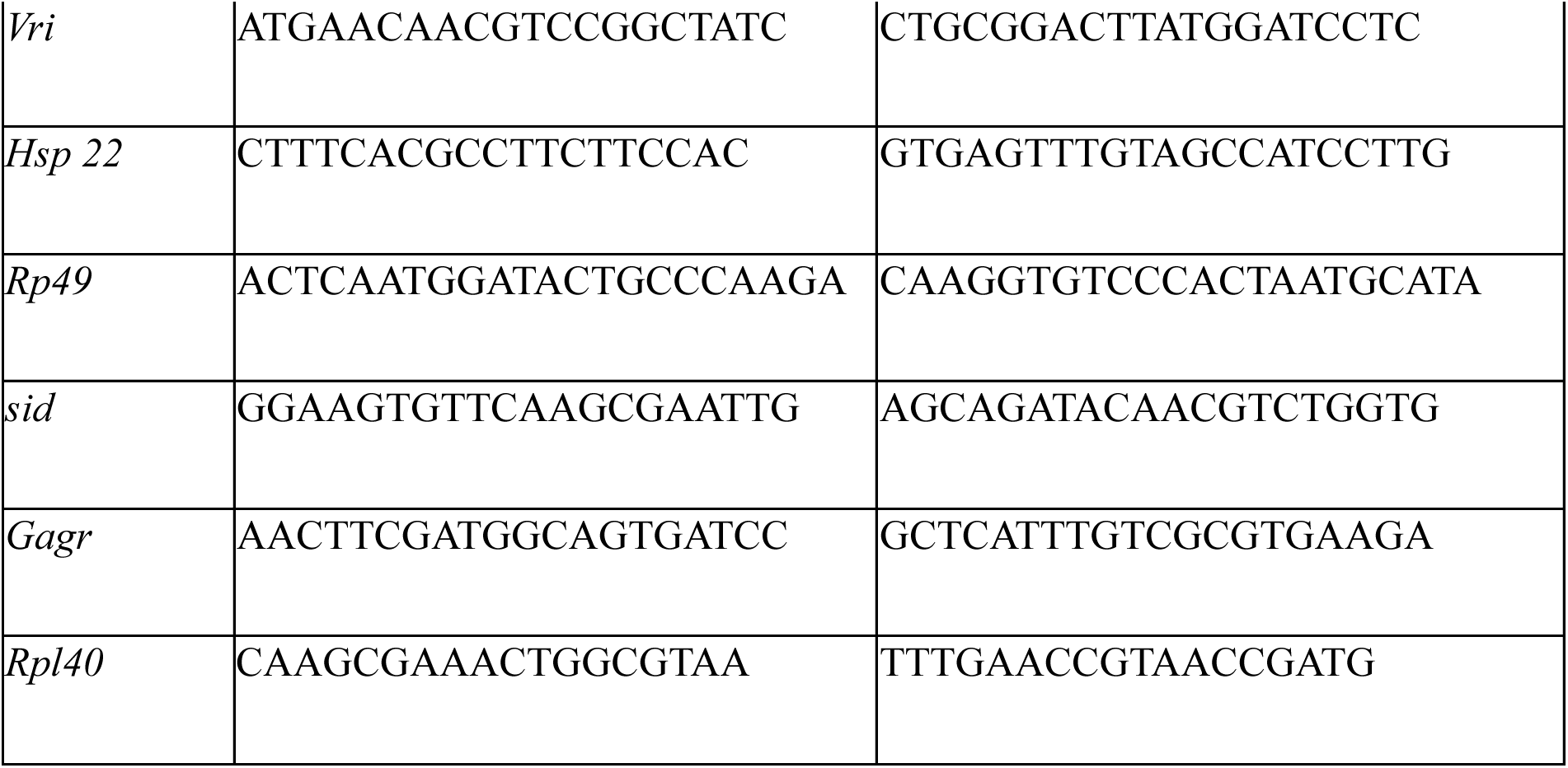
List of genes and primers used for gene expression analysis.

**Table 1.S2:**
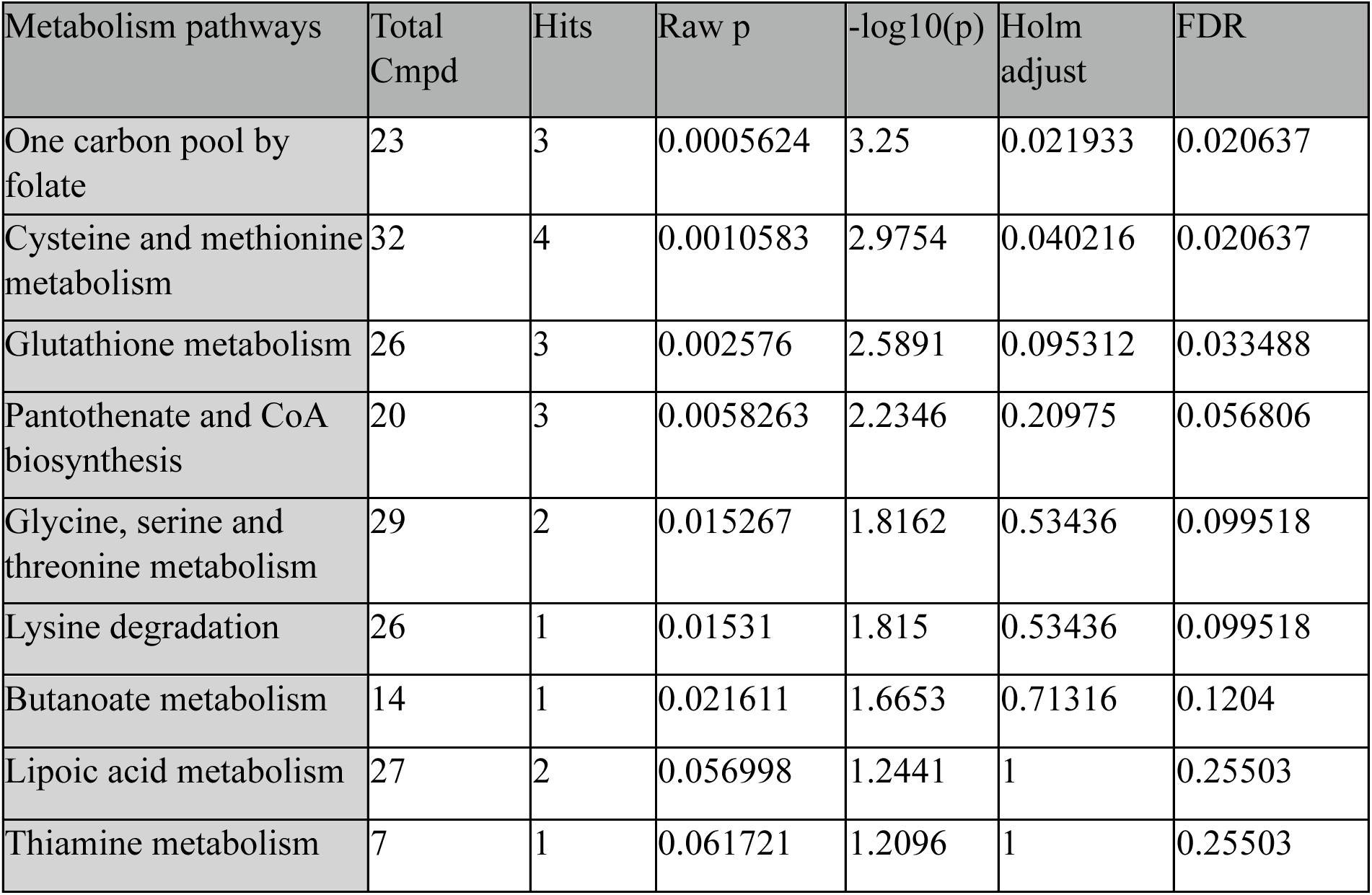
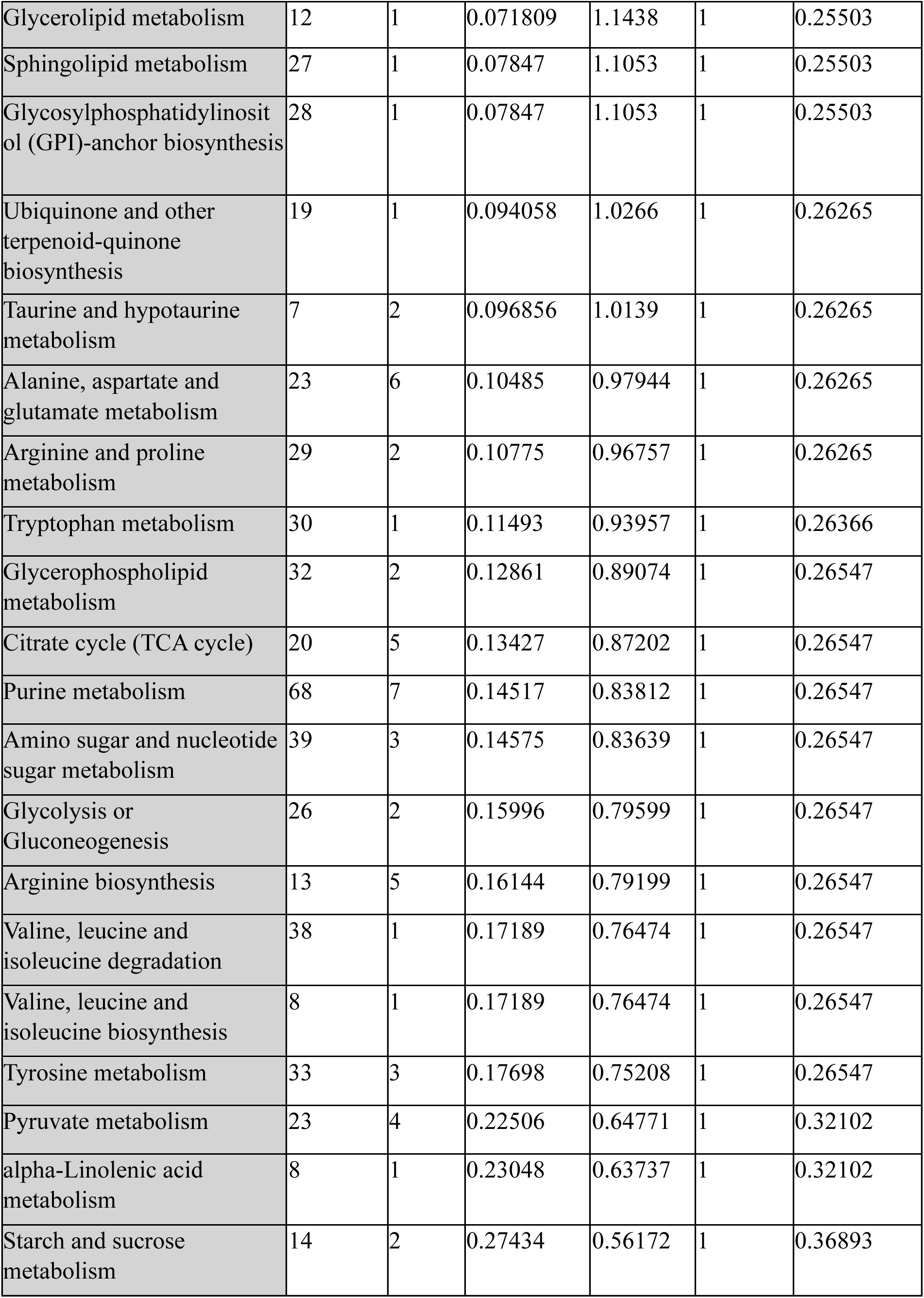
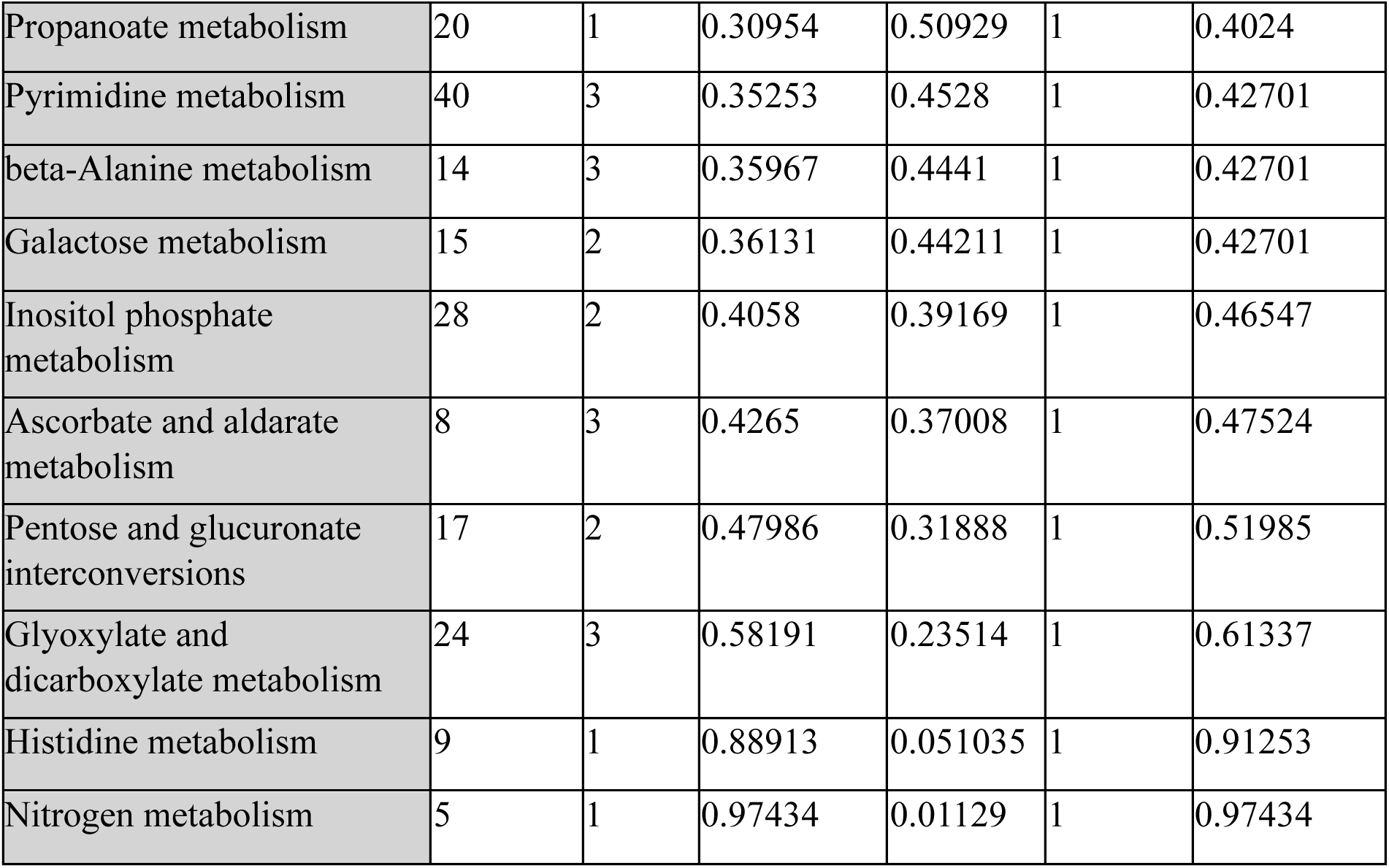
Pathways enriched in PTEF post-exercise, and the total compounds contributing to these pathways.

